# From Consensus to Individual Differences: Typicality Links Brain and Behavior Across Naturalistic Contexts

**DOI:** 10.64898/2026.08.14.744685

**Authors:** Michal Zamberg-Elad, Ido Har-Shalom, Aviv Jarbi, Meytal Wilf, Michal Ramot

## Abstract

When people experience the same naturalistic events, their brains and behavior converge on remarkably similar responses. This shared response is widely used to identify what people have in common, but whether the consensus itself carries meaning has remained largely unexplored. Here we propose that the group consensus is not merely an average, but an approximation of a shared solution to the computation being performed, such that an individual’s proximity to it, their typicality, indexes how well they perform that computation. Across behavioral tasks spanning social and non-social cognition, typicality was a stable characteristic of the individual, yet specific to the computation engaged rather than a general tendency to conform. Neural typicality during movie viewing showed the same organization, and was linked to behavior most strongly where the relevant computation was engaged: typicality in face-selective cortex was more strongly related to behavior on tasks involving faces than on matched tasks without faces, particularly during movies containing faces. Critically, where ground truth was available, proximity to the consensus predicted objective performance: neural typicality measured during passive movie viewing predicted accuracy on independent motion-prediction and face-recognition tasks. These findings suggest that, when individuals share objectives and constraints, the group consensus can serve as an empirical benchmark of successful processing, offering a principled way to measure individual differences in complex behaviors for which no correct answer can be defined.

**Significance Statement:** Many complex human behaviors, such as interpreting another person’s emotions, have no single “correct” response, making them difficult to quantify using traditional measures. We propose that the response people converge on when experiencing the same events is not merely an average, but an approximation of a shared solution, so that how closely an individual approaches it, their typicality, can serve as a measure of performance. We show that typicality is a stable characteristic of the individual, specific to the type of information being processed, and linked to the brain systems supporting it. Where correct answers exist, people whose brain responses during movie viewing were more typical performed better on independent tasks. These findings provide a new way to study individual differences in complex, real-world behavior and suggest that shared responses may, under appropriate conditions, reveal how effectively information is being processed.

## Introduction

Understanding the link between neural activity and behavior remains one of the central challenges of cognitive neuroscience. While the ability of non-invasive neuroimaging to elucidate whole-brain dynamics continues to improve, establishing robust brain-behavior correlations is ultimately constrained by our ability to measure both our behavior of interest and neural activity in a manner that captures stable, trait-like variance rather than transient state fluctuations. This challenge becomes especially critical when neural and behavioral data are acquired at different time points or under different task conditions. Yet for many complex, naturalistic behaviors, defining such stable and objective measures is difficult.

Many of the behaviors we most wish to understand are complex, involve multiple underlying processes, and lack a clear external ground truth. Natural behaviors are rich and dynamic, and difficult to reduce to a single score. In social interactions, for instance, emotional signals are rarely conveyed as categorical, unambiguous states. What constitutes the “correct” interpretation of an emotionally complex narrative? How should empathic accuracy be quantified when the emotion experienced by the speaker, their intended message, and the listener’s perception may all diverge? (Jospe et al. 2020) Even gaze behavior during naturalistic viewing, although highly constrained on average by cinematic structure (Hasson et al. 2008), exhibits substantial and meaningful inter-individual variability (Dorr et al. 2010; Ramot et al. 2020). Traditional scoring approaches either rely on externally imposed labels, or reduce complex behavior to simplified proxies, often obscuring the structure that makes natural behavior informative in the first place.

We propose the concept of *typicality* as an organizing principle for addressing these challenges. Typicality reframes individual differences in relation to a stable group consensus: when independent observers repeatedly produce similar responses to the same stimulus, their shared response provides an empirical reference against which individual variation can be measured, even when no objective ground truth can be specified a priori. Importantly, this consensus may provide more than a descriptive average. When individuals are exposed to the same information and operate under similar goals and constraints, their responses contain both shared and idiosyncratic components (Chamberlain et al. 2024). To the extent that idiosyncratic variation is not systematically aligned across individuals, averaging attenuates this variation while preserving information that is consistently shared across observers. This logic is related to the wisdom-of-crowds principle, in which aggregation of partially independent errors can produce a more accurate estimate than any single judgment (Galton 1907; Surowiecki 2004) and is central to intersubject approaches in naturalistic neuroscience, which leverage shared stimulus-locked responses to identify information consistently represented across individuals (Hasson et al. 2004; Nastase et al. 2019). Typicality then quantifies an individual’s deviation from this empirically estimated shared response, providing a principled measure of individual variation in both behavior and neural activity.

Naturalistic cognition provides a particularly strong basis for defining such a consensus. Naturalistic stimuli evoke robust intersubject correlations (ISC) across widespread cortical regions, spanning sensory, associative, and default-mode networks (Hasson et al. 2004, 2010; Chen et al. 2017; Nguyen et al. 2019). The degree of neural similarity between individuals has been linked to shared memory (Chen et al. 2017), shared interpretation (Nguyen et al. 2019), and even social proximity (Parkinson et al. 2018), demonstrating that convergence can reflect meaningful commonalities in processing rather than merely common sensory input (Chamberlain et al. 2024). These methods provide powerful measures of shared neural processing. Inter-subject representational similarity analyses have further linked neural similarity to performance, showing that high performers on cognitive tasks tend to be more synchronized with one another, whereas low performers are idiosyncratic in their own ways (An “Anna Karenina” pattern; Finn et al. 2020; Chamberlain et al. 2024). Such findings indicate that similarity between individuals is not behaviorally neutral. They remain, however, grounded in how similarly individuals respond to one another, and leave open whether the consensus itself carries meaning. Because substantial individual variability remains around these shared responses this question can be addressed directly, by asking whether an individual’s proximity to the consensus reflects more effective cognitive processing.

Our previous work took a first step in this direction, using variability around the shared response to link behavioral and neural measures of typicality during social perception. Individual differences in gaze typicality measured while participants viewed one set of social movies predicted their neural typicality measured during an independent movie, revealing a distributed network associated with social orienting (Ramot et al. 2020). This brain-behavior relationship generalized across different stimuli and measurement contexts, inside and outside the MRI scanner, demonstrating that an individual’s distance from the group response can capture a stable and behaviorally meaningful characteristic rather than a stimulus-specific fluctuation. Whether this relationship generalizes beyond social gaze, and more fundamentally what proximity to the shared response signifies, remains unknown.

These observations raise a broader set of questions about the relationship between shared responses and individual differences. First, is typicality generally a reliable characteristic of the individual across behavioral and neural domains, or was the stability we previously observed specific to social gaze? Second, does an individual’s typicality ranking reflect their global tendency to respond like others, perhaps because of attentiveness, compliance, or a general propensity toward consensus, or is it sensitive to the particular computation being performed? If the latter is true, typicality should generalize strongly across independent stimuli engaging similar processes, but less strongly across conditions engaging different processes. Correspondingly, behavioral typicality on a given task should relate most strongly to neural typicality in systems engaged by the relevant computation, and during stimuli that engage it. Most fundamentally, what does proximity to the consensus signify? Does the shared response simply provide a useful reference for quantifying individual differences, or does an individual’s proximity to it carry information about how successfully the underlying computation is being performed?

Here, we systematically examine these questions across both behavioral and neural domains. Across multiple tasks spanning both social and non-social contexts, we demonstrate that behavioral typicality is stable within individuals, yet sensitive to content and task demands. In parallel, we show that neural typicality during naturalistic viewing is stable across independent stimuli, while also remaining sensitive to stimulus content. Crucially, we find that behavioral typicality is related to neural typicality across multiple domains, with different behavioral measures mapping onto neural systems relevant to the corresponding computations.

Finally, we ask whether the consensus itself carries additional information about successful processing. Agreement with the group is not inherently equivalent to better performance. However, in tasks with independently defined correct responses, we find that proximity to the behavioral consensus is strongly associated with objective performance, and that neural typicality measured during naturalistic movie viewing predicts individual differences in performance. These findings suggest that, under conditions in which individuals share similar objectives and constraints, the consensus may approximate an effective solution to the computation being performed, and typicality may index how closely an individual approaches it. In this sense, consensus provides not only a reference for quantifying individual differences but also a potential means of estimating successful processing when objective ground truth cannot be defined.

## Results

### Behavioral Typicality Is a Reliable Trait Sensitive to Category-Specific Processing

A central aim of this study was to test whether typicality can serve not only as a reliable individual measure, but as a metric sensitive to the specific processes engaged by a stimulus. We therefore designed a behavioral battery spanning a broad range of cognitive functions, including both social and non-social tasks across several modalities: naturalistic movie viewing with eye tracking of social / animal / object videos; intensity ratings of auditory emotional narratives / non-social complex soundscapes; biological / non-biological motion prediction; and recognition memory (Cambridge Face / Cambridge Car Memory Test) (Figure 1; see Methods). Each task pairs social/non-social conditions that share low-level demands but are thought to recruit partially distinct mechanisms due to the different nature of the stimuli in each condition. For example, attention during naturalistic viewing is driven by shared cues such as salience, motion, and contrast across all categories, but social scenes additionally engage social-specific orienting. We reasoned that if typicality reflects only a generic disposition, such as a consistent rating style, general attentiveness, or a global tendency toward consensus, it should be expressed uniformly across these conditions. If instead it is sensitive to the specific computation engaged, its structure should differ across conditions (but remain stable across different stimuli within condition) in a manner that mirrors the known similarities and differences between them.

**Figure 1.**
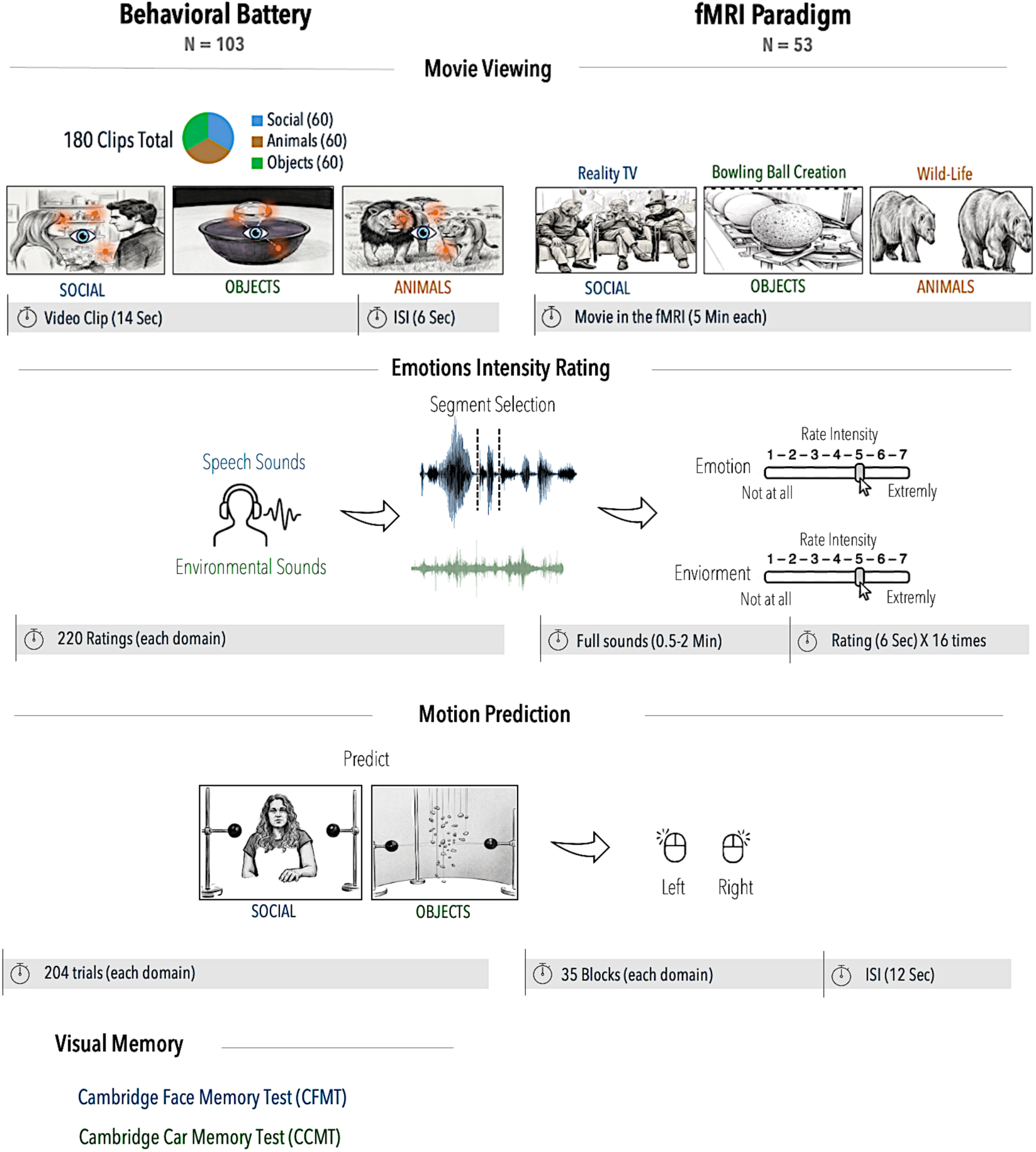
Overview of the experimental design and behavioral paradigms. Behavioral and fMRI paradigms used to assess neural and behavioral typicality and scores. Participants in the behavioral cohort (*N* = 103) completed **(a)** eye-tracking during viewing of 180 short movies (60 social, 60 object, and 60 animal), **(b)** emotion- and sound-intensity ratings, **(c)** biological and non-biological motion prediction tasks, and **(d)** recognition memory assessments using the Cambridge Face Memory Test (CFMT) and Cambridge Car Memory Test (CCMT). Participants in the fMRI cohort (*N* = 53) viewed three naturalistic movies representing social, object, and animal content while BOLD activity was recorded. Neural typicality was computed by correlating each participant’s voxelwise time course with the leave-one-out group average for each movie. Two participants were excluded from each movie (missing data or low engagement, defined as mean neural typicality more than 2 SD below the sample mean; see Methods), yielding a final sample of *N* = 51 per movie. They also conducted the emotion ratings and motion prediction tasks inside the fMRI.

Because typicality is defined as an individual’s deviation from the group-consensus response, it relies on the existence of a stable consensus in the first place. We therefore began by confirming convergence to a stable mean. For the movie watching paradigm, participants viewed 180 short movie clips, 60 in each category (see Methods). A split-half reliability analysis confirmed that in 161 out of 180 movies, gaze trajectories converged to a very stable average (0.8 < r < 0.998; see Methods). We further chose a subset of movies (45 social, 45 animals and 43 objects) that match on the mean and variance of the typicality values. These were the movies included in all further analyses.

Having established that on average participants converge to a stable consensus, we next tested whether an individual’s distance from this shared response was stable across stimuli and across domains (social/non-social). We quantified each participant’s gaze typicality as the distance between their moment-to-moment gaze trajectory and the leave-one-out group-average trajectory for each of the short movie clips they viewed. We then compared the consistency of individual gaze typicality scores across different subsets of videos within each stimulus category to their consistency across categories (Figure 2a). Within all three categories, gaze typicality was highly reliable across independent subsets of movies (social: r = 0.85, SD = 0.02, 95% CI [0.80, 0.89]; animal: r = 0.76, SD = 0.04, 95% CI [0.68, 0.82]; object: r = 0.79, SD = 0.03, 95% CI [0.72, 0.85]), indicating that typicality captures stable individual differences rather than transient fluctuations or noise. Because the movie clips were presented across three separate sessions, this reliability reflects consistency of gaze typicality across sessions and stimuli, not only within a single sitting. This within-category reliability, particularly for the social category, replicates our previous finding that gaze typicality is a stable individual characteristic that generalizes beyond the specific stimulus (Ramot et al. 2020).

**Figure 2.**
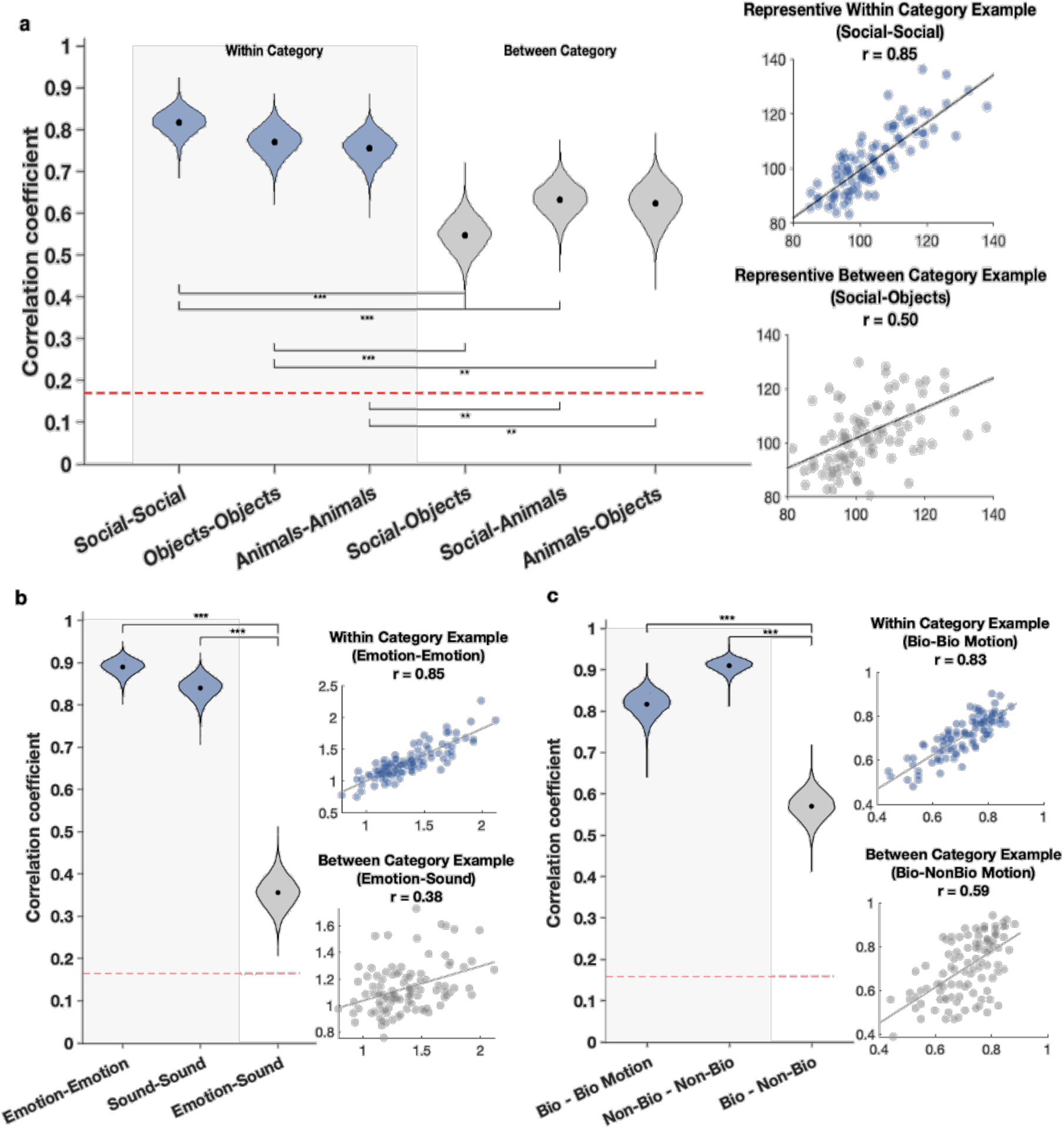
Split-half reliability of typicality across behavioral categories. Correlation distributions were obtained by randomly splitting stimuli within each category into two independent halves across 10,000 iterations and computing the correlation between subject-level estimates derived from each half. Histograms show the resulting distribution of correlation coefficients, for within-category split-half correlations (blue) or cross-category correlations (gray), as indicated in each panel. The red dashed line marks the 95th percentile of the null distribution generated by randomly permuting subject identities. Representative scatterplots from a single iteration illustrate individual split-half or cross-category relationships. **(a)** Movie-viewing typicality. Within-category correlations were consistently higher than cross-category correlations (all p ≤ .036), with mean correlations of 0.85 (Social), 0.79 (Objects) and 0.76 (Animals), compared with 0.56 (Social–Objects), 0.63 (Social–Animals) and 0.65 (Animals–Objects). **(b)** Emotion- and sound-rating typicality. Emotion-(mean r = 0.89) and sound-rating (mean r = 0.84) typicality exhibited substantially higher reliability than cross-category correlations (mean r = 0.36; both p < .0001). **(c)** Motion-prediction accuracy. Within-category reliability was high (biological: mean r = 0.82; non-biological: mean r = 0.91), whereas cross-category correlations were lower (mean r = 0.57; both p < .0001).

Though still relatively high and well above chance, between-category correlations were significantly lower than within-category reliability (social-object: r = 0.56, 95% CI [0.49, 0.64]; social-animal: r = 0.63, 95% CI [0.56, 0.70]; animal-object: r = 0.65, 95% CI [0.57, 0.72], Figure 2a), as expected given the low-level attentional mechanisms shared across all naturalistic viewing. Comparing within- and between-category correlations across the 10,000 random splits, within-category reliability significantly exceeded between-category correlations for social movies (vs. social-objects: Δ = 0.288, 95% CI [0.204, 0.369], p < .0001; vs. social-animals: Δ = 0.217, 95% CI [0.137, 0.299], p < .0001), object movies (vs. social-objects: Δ = 0.227, 95% CI [0.135, 0.315], p < .0001; vs. animals-objects: Δ = 0.137, 95% CI [0.046, 0.227], p = .0026), and animal movies (vs. social-animals: Δ = 0.122, 95% CI [0.022, 0.213], p = .019; vs. animals-objects: Δ = 0.102, 95% CI [0.008, 0.195], p = .036). Importantly, the magnitude of these differences varied across categories. The largest within-versus between-category gaps were found for social movies. Social viewing was the most distinct, separated from both non-social categories, whereas animal and object movies were more similar to one another (Figure 2). Across splits, the within-versus-between gap was larger for social movies than for the animal-object comparison (Δ = 0.115, 95% CI [0.011, 0.221], p = .028).

To test whether this organization depended on our a priori category labels, we performed an unsupervised analysis in which movies were clustered solely by their across-individual gaze-typicality profiles. The social movies recovered a single, high-purity cluster (purity = 0.911, permutation p < 0.0001), whereas the animal and object clusters had lower purity (animals purity = 0.44, objects purity = 0.63) (Supplementary Figure S1). Thus, the distinction of social viewing emerged independently of the predefined stimulus categories.

We also observed a stable group consensus and stable individual distances from the consensus in the auditory task. Participants listened to short audio clips consisting of either conversational speech or environmental (non-social) sounds and rated the intensity of brief segments extracted from each clip on a 1–7 scale: emotional intensity for speech and acoustic intensity for environmental sounds (see Methods), for which there is no objectively correct response. Participants nevertheless converged on highly reliable consensus ratings for both emotional speech (mean split-half *r* = 0.97, SD = 0.004, 95% CI [0.96, 0.98], Figure 2b) and environmental sounds (mean split-half *r* = 0.99, SD = 0.002, 95% CI [0.98, 0.99]), indicating that participants showed consistent agreement in both domains. Individual emotional-intensity typicality scores (mean *r* = 0.89, SD = 0.02, 95% CI [0.85, 0.92]) and sound-intensity typicality scores (mean *r* = 0.84, SD = 0.02, 95% CI [0.79, 0.88]) were highly reliable within condition, yet substantially less reliable across conditions (emotion–sound: mean *r* = 0.36, SD = 0.04, 95% CI [0.28, 0.43]; both within-versus-between comparisons, *p* < 0.0001). These findings indicate that an individual’s rating typicality does not reflect a general response style applied uniformly across tasks, but rather process-specific sensitivity that distinguishes social (empathic) from non-social (perceptual) judgments.

The motion-prediction task showed a similar pattern, while also allowing us to compare typicality with objective performance. Participants viewed clips truncated shortly after motion onset, depicting either a human agent (social, biological motion) or an inanimate object (non-social, non-biological motion) about to move left or right, and predicted the upcoming direction of movement from these minimal cues (see Methods). Because an objective ground truth exists, we quantified both prediction accuracy and prediction typicality (deviation from the group response). As with previous tasks, participants converged on highly reliable consensus responses in both conditions (see Methods, Supplementary Figure S2c). Both accuracy and typicality were also highly reliable within category (accuracy: biological motion mean r = 0.82, SD = 0.03, 95% CI [0.75, 0.87]) and non-biological motion (mean r = 0.91, SD = 0.02, 95% CI [0.88, 0.94]; typicality: biological motion mean r = 0.83, SD = 0.03, 95% CI [0.77, 0.88], Figure 2c) and non-biological motion (mean r = 0.91, SD = 0.02, 95% CI [0.88, 0.94]). Across category correlations between biological and non-biological prediction accuracy were significantly lower than within category for both accuracy and typicality (accuracy: mean r = 0.57, SD = 0.03, 95% CI [0.50, 0.63]; typicality: mean r = 0.55, SD = 0.04, 95% CI [0.48, 0.61]; p < 0.0001 for both compared to within category). Importantly, prediction typicality was strongly associated with objective task performance, with more typical responses corresponding to higher prediction accuracy for both biological motion (r = −0.87, p < 0.001) and non-biological motion (r = −0.96, p < 0.001). Thus, when an objective ground truth exists, greater agreement with the group is associated with more accurate performance.

Finally, behavioral typicality scores were largely independent across tasks. After correcting for multiple comparisons (Benjamini–Hochberg false discovery rate), only two of the 30 cross-task comparisons remained significant: between the Cambridge Car Memory Test (CCMT) and both auditory rating typicality (emotion ratings: r = −0.31, FDR-adjusted p = 0.022; sound ratings: r = −0.31, FDR-adjusted p = 0.022). No other cross-task associations survived correction (all FDR-adjusted p > 0.05). Together, these findings indicate that behavioral typicality is not a single global disposition toward consensus. Rather, it is a reliable measure that remains sensitive to the type of information being processed.

### Neural Typicality Reveals Both General and Content-Specific Processing Networks

Having established that behavioral typicality is stable within individuals while remaining sensitive to stimulus content, we next examined whether neural typicality during naturalistic viewing displays similar properties.

Participants viewed three 5-minute movies in the 7T fMRI scanner: one wildlife movie, one objects movie (the production process of a bowling ball, no humans in the video), and one social movie, which was completely distinct from the short movie clips used to calculate gaze typicality. Neural typicality was defined, for each participant and voxel, as the correlation between that voxel’s time course and the average time course of all other participants at that voxel (see Methods). We then examined how neural typicality varied across movie categories.

Average neural typicality maps revealed widespread synchronization across participants for all three movie categories (Figure 3a–c). All three maps showed extensive effects well above their respective null-derived thresholds (see Methods). Because significant typicality extended across large portions of the cortex, the overlap map (Figure 3d) is restricted to the strongest effects within each movie to facilitate visualization of shared and category-specific regions.

**Figure 3.**
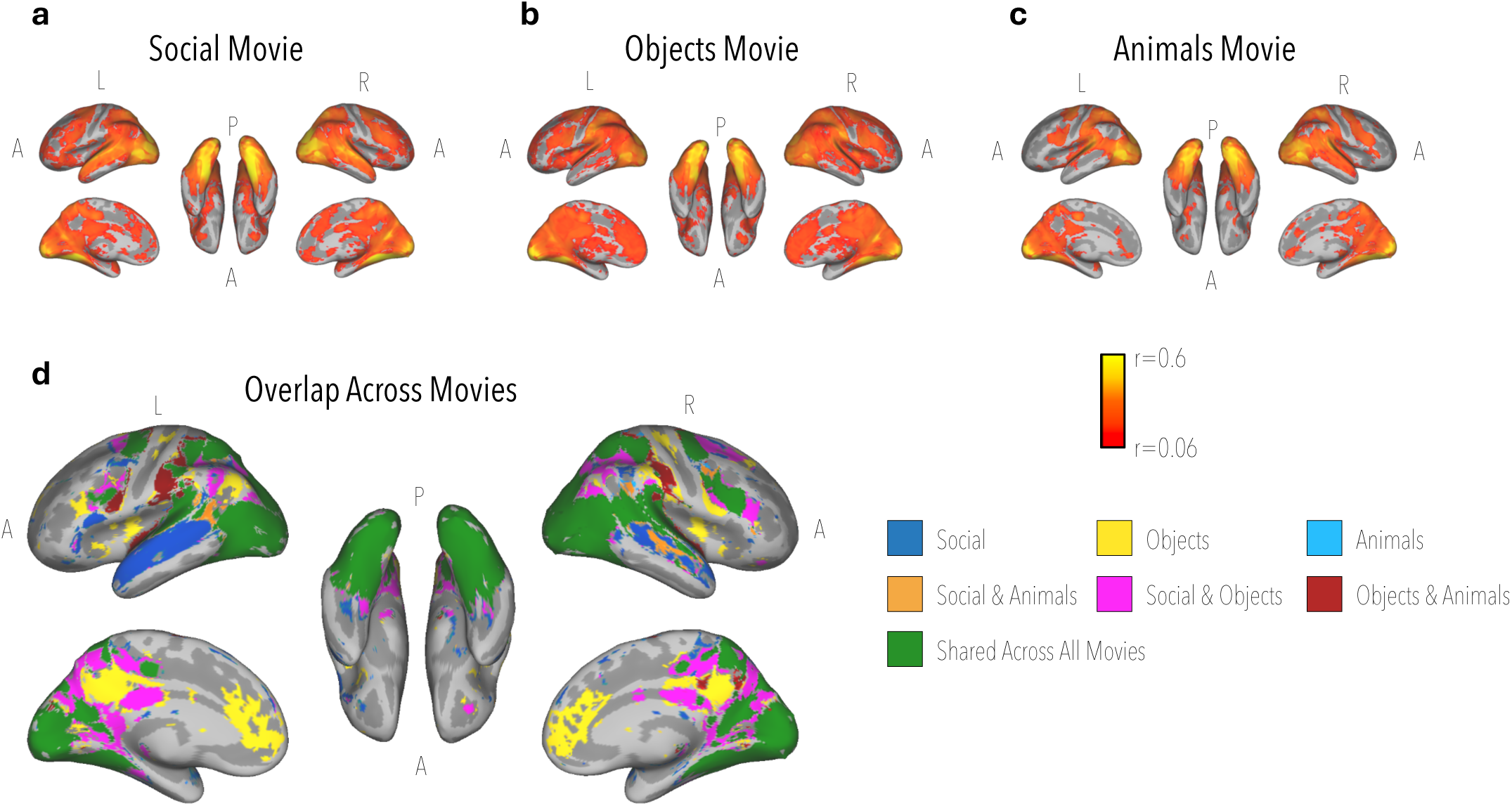
Neural typicality maps across movie categories. Average neural typicality during social (**a**), object (**b**), and animal (**c**) movie viewing. Visual and auditory cortices exhibited robust typical responses across all movie categories. Maps in (**a–c**) are thresholded at each movie category’s own null-derived threshold, obtained via a time-reversal permutation null model (see Methods): |r| > 0.076 for social (**a**), |r| > 0.064 for object (**b**), and |r| > 0.099 for animal (**c**). The overlap map (**d**) highlights regions uniquely associated with each category, pairwise overlaps, and regions exhibiting typical responses across all movie types; for visualization, this map retains the strongest 70% of voxels (by effect size) within the null-thresholded maps.

Visual and auditory cortices exhibited strong typical responses regardless of movie content, indicating that all stimuli elicited highly consistent sensory processing across viewers. Overlap analyses revealed a common sensory network shared across social, animal, and object movies (Figure 3d). Beyond these shared sensory regions, however, marked differences emerged between movie categories in the regions exhibiting the strongest synchronization across participants. Social movies elicited extensive neural typicality within regions classically associated with language and social cognition, including the superior temporal sulcus (STS), medial prefrontal cortex, temporoparietal regions, and frontal association cortex (Lahnakoski et al. 2012; Hasson et al. 2004; Nastase et al. 2019, 2020; Fletcher 1995). Animal movies recruited partially overlapping networks, particularly within the right STS, while exhibiting less extensive recruitment of higher-order social regions. In contrast, object movies showed stronger involvement of medial visual regions and object-selective cortex. Object and animal movies showed more limited pairwise overlap, concentrated primarily in dorsal frontal and parietal association regions implicated in visuospatial attention and the processing of dynamic visual information.

Several frontal regions, especially in right dorsolateral prefrontal cortex as well as parts of the intraparietal sulcus, exhibited strong typical responses across all movie categories, suggesting the existence of domain-general executive control and attention systems involved in naturalistic processing. However, the dominant contribution of social brain regions during social movie viewing indicates that neural typicality reflects far more than shared attention or engagement. Instead, typicality appears sensitive to the specific cognitive demands imposed by different types of naturalistic stimuli.

### Individual Neural Typicality Combines Stability Across Stimuli with Stimulus-Specific Organization

We next asked whether neural typicality reflects a stable characteristic of the individual or whether it is entirely determined by stimulus content. We compared individual neural typicality both within movies, using independent halves of the same movie, and across different movie categories (Supplementary Figure S3). Across large portions of the cortex, the relative neural typicality of individual participants was preserved across both independent halves of the same movie and different movies.

Importantly, however, this stability depended on movie content. Across-movie stability was strongest in regions engaged by both movies being compared, whereas regions preferentially engaged by a particular movie showed stable neural typicality primarily within that movie (for instance, STS reliably captured individual neural typicality profiles mostly within the social movie context). Thus, as in the behavioral measures, neural typicality contains both a component that is stable across stimuli and a stimulus-dependent component, reflecting an interaction between stable individual tendencies and the computations engaged by the stimulus. Because the movies were distributed across two scanning sessions, between-movie comparisons for some participants spanned separate sessions, providing partial evidence that this stability extends beyond a single scanning session.

### Social Gaze Typicality Correlates to Neural Typicality Within Social Brain Networks

Having established the reliability of both behavioral and neural typicality, we next examined whether behavioral typicality correlates with neural typicality. Social gaze typicality measured outside the scanner was strongly associated with neural typicality during social movie viewing (Figure 4a). Significant correlations were observed in language areas in the superior temporal sulcus and inferior frontal cortex, as well as in portions of the default mode network (also see Supplementary Figure S7a for semantic decoding and network correspondence analysis). Results survived whole-brain cluster correction (see Methods), with a peak correlation of r = −0.64; this relationship also survived a conservative family-wise correction (see Methods). Negative correlations are expected, as gaze typicality is a distance measure (higher values indicating reduced typicality), whereas neural typicality is a correlation metric, where higher values indicate greater typicality. No voxels showed significant correlations in the opposite direction, meaning that more typical gaze always corresponded to more typical neural responses.

**Figure 4.**
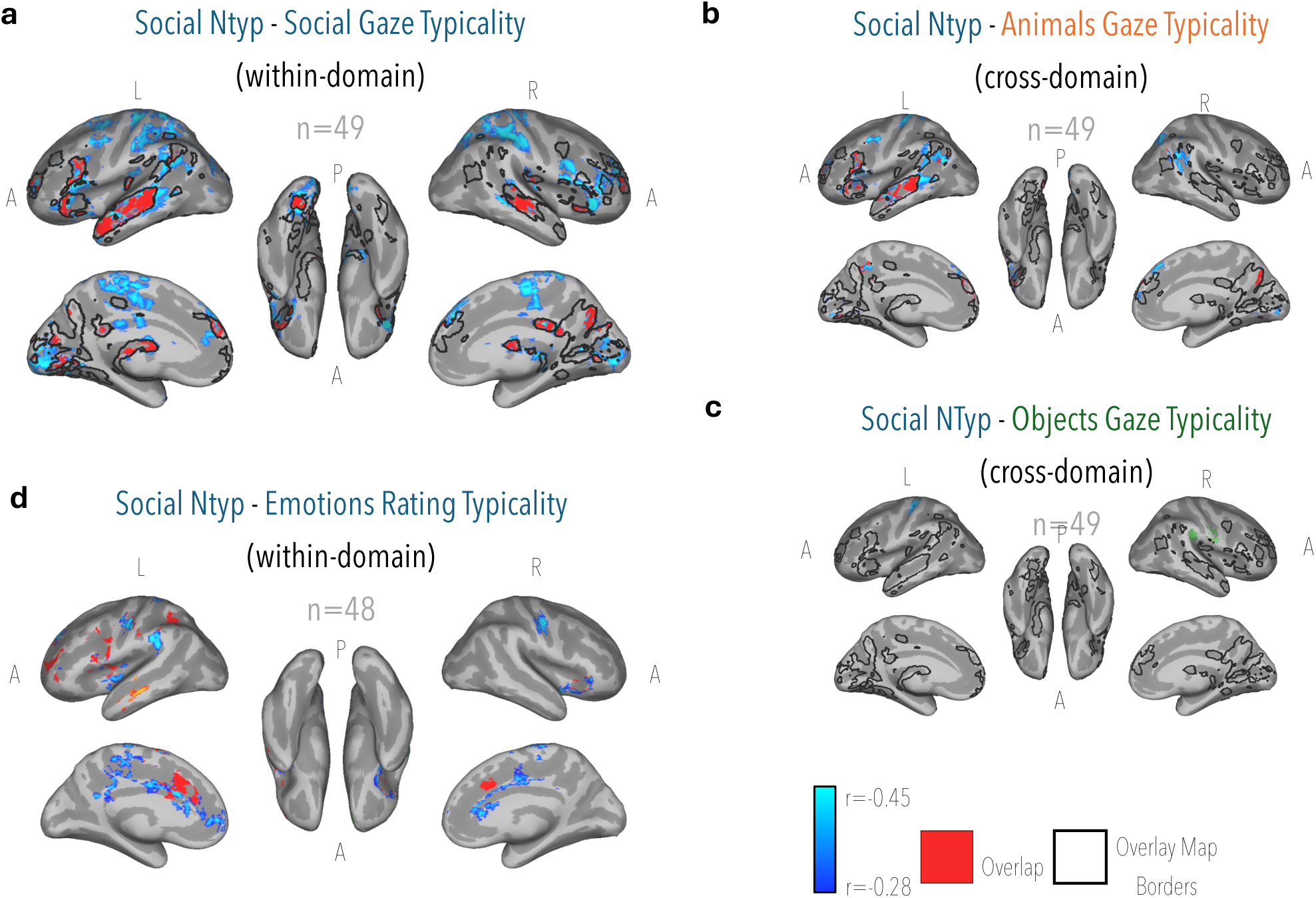
Correlations between social neural typicality and behavioral typicality across content categories. Voxelwise correlation maps showing the relationship between neural typicality (NTyp) during social movie viewing and gaze typicality derived from social **(a)**, animal **(b)**, and object **(c)** movies in the behavioral paradigm. Maps display only voxels surviving cluster-correction. Each analysis was thresholded using its own cluster-size threshold. **(a)** The black contour outlines the region identified by Ramot et al. (2020) and illustrates its overlap with the current findings (overlap in red). **(d)** Voxelwise correlation maps showing the relationship between social neural typicality (NTyp) and emotion-rating typicality derived from the behavioral battery. The red regions indicate the overlap between the map and the activation map from the emotion-rating GLM.

Importantly, this brain–behavior map reproduced the social orienting network we previously identified through the same typicality logic (Ramot et al. 2020, Figure 4; *black* contours*, red* overlap). This replication was obtained in an independent cohort, using a different scanner and different movie stimuli in a different language. Moreover, behavioral gaze typicality and neural typicality were again derived from independent sets of movies, demonstrating that the relationship generalizes beyond the particular stimuli used to measure either behavior or neural responses.

Social movie neural typicality was also associated with gaze typicality measured during animal movies, although these effects were less spatially extensive (Figure 4b). The association survived whole-brain cluster correction (see Methods), with a peak correlation of r = −0.60. In contrast, gaze typicality measured during object movies showed only limited associations with neural typicality during the social movie (Figure 4c, peak correlation r = - 0.53, whole-brain cluster corrected). The overlay of these maps is shown in Supplementary Figure S4a-b. Neural typicality during the object movie produced few relationships with gaze typicality, even for object gaze, and these were inconsistent in direction (Supplementary Figure S5).. Thus, neural typicality tracked gaze behavior most strongly in contexts in which gaze was socially informative, mirroring the category structure observed behaviorally rather than reflecting a generic correspondence between typicality measures.

### Emotional Interpretation Correlates to Neural Typicality Beyond Gaze Behavior

We next asked whether the relationship between behavioral and neural typicality extends to other aspects of social cognition beyond gaze behavior. Emotion-rating typicality was associated with neural typicality within medial frontal cortex, insula, and other regions implicated in affective and social processing (Figure 4d). The association survived whole-brain cluster correction (see Methods), with a peak correlation of r = −0.60. Notably, these regions overlapped substantially with areas identified by the task-based emotion analyses, especially in frontal regions (Figure 4d, red overlap), providing independent evidence that the relationship was localized to regions engaged during emotional processing (also see Supplementary Figure S7b for semantic analysis).

In contrast, sound-intensity typicality showed few significant associations with neural typicality that survived correction for multiple comparisons. Moreover, there was almost no overlap between the neural correlates of emotion-rating and sound-intensity typicality (Supplementary Figure S4d), despite the two measures sharing the same rating structure. Thus, the relationship between emotion-rating typicality and neural typicality cannot be explained simply by a general tendency to produce typical intensity ratings.

### Neural Typicality Predicts Performance on Motion Prediction Tasks

The previous analyses demonstrated that neural typicality was associated with behavioral measures defined relative to the group consensus. We next asked whether neural typicality predicts performance on tasks with objectively defined correct answers, beginning with motion prediction, for which we have shown a high correspondence between behavioral typicality and accuracy. Neural typicality during social movie viewing was positively associated with biological motion prediction accuracy (51,560 voxels after whole-brain cluster correction; peak r = 0.67; N = 48; Figure 5a; see Supplementary Figure S7c for semantic analysis), predominantly in visual cortex, lateral temporal cortex including the superior and middle temporal gyri, and the fusiform gyri, as well as in superior parietal cortex, supplementary motor area and inferior frontal cortex; this relationship also survived a conservative family-wise correction (see Methods). Neural typicality during object movie viewing was associated with non-biological motion prediction accuracy (26,734 voxels; peak r = 0.63; N = 48; Figure 5b), predominantly in medial parietal cortex (precuneus and posterior cingulate), the angular gyri and medial and dorsal frontal cortex, as well as the middle temporal gyrus and hippocampus; this relationship likewise survived the family-wise correction. The two maps showed little spatial overlap (2,428 shared voxels, 5% and 9% of the respective maps; Supplementary Figure S4f), suggesting that neural typicality was associated with biological and non-biological motion prediction through largely distinct neural systems. Descriptively, biological motion prediction was related to typicality more extensively during the social than during the object movie (51,560 vs 17,542 voxels), and non-biological motion prediction more extensively during the object than during the social movie (26,734 vs 16,033 voxels; Figure 5c). However, a direct test of this task × movie crossover within a common mask, which controls for global differences between participants by permuting movie labels within each participant, was not significant (p = .18; within independent motion-task masks, p = .06 and .10), nor was the preference of biological motion prediction for the social movie on its own (p = .20; p = .09 within its task mask). Correlations with both tasks were positive during both movies, suggesting that part of the relationship reflects factors shared across movies and tasks. Because performance in both tasks was quantified against objectively correct responses, these findings extend the relationship between neural typicality and behavior beyond consensus-based measures: individuals whose neural responses during naturalistic viewing were more typical tended to predict motion more accurately.

**Figure 5.**
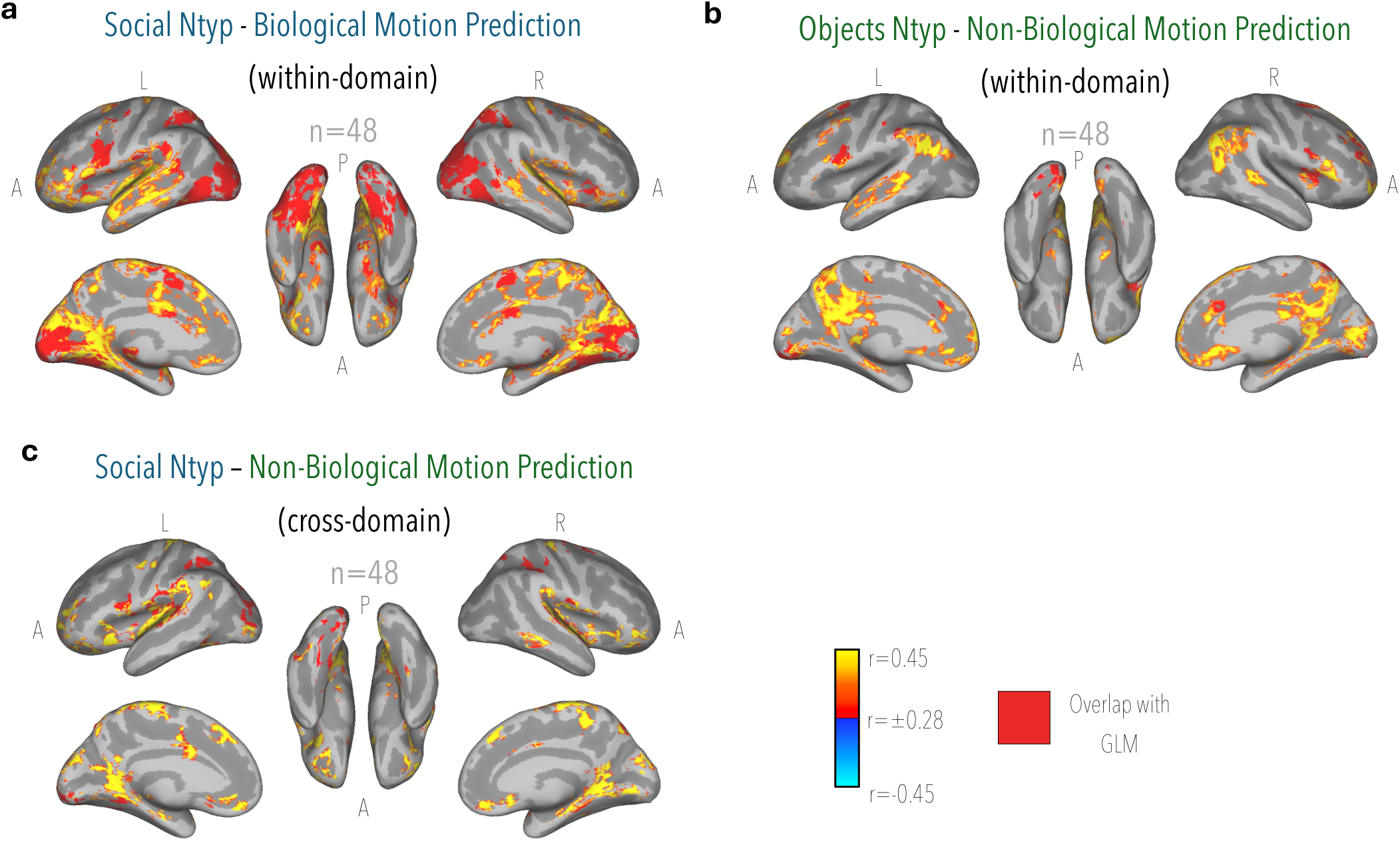
Neural typicality predicts biological motion prediction. Voxelwise correlation maps showing the relationship between neural typicality (NTyp) movie viewing and behavioral performance. **(a)** Brain regions in which social neural typicality correlates with biological motion prediction accuracy. In red is the overlap between the correlation to the task GLM, where the biological motion is significant compared to rest. **(b)** Brain regions in which object neural typicality correlates with non-biological motion prediction accuracy. In red is the overlap between the correlation to the task GLM, where the non-biological motion is significant compared to rest. **(c)** Brain regions in which social neural typicality correlated with non-biological motion prediction accuracy. In red is the overlap between the correlation to the task GLM, where the non-biological motion is significant compared to rest. Maps display only voxels that survive cluster-correction (see Methods).

### Neural Typicality Predicts Face Recognition Performance

Having established that neural typicality predicts objective performance in motion prediction, we next asked whether this relationship extends to face-recognition ability, measured using the Cambridge Face Memory Test (CFMT). As in the motion-prediction task, the CFMT also allowed us to directly compare behavioral typicality with objective accuracy. The two measures were very strongly correlated (r = −0.94, p < 0.0001), providing a second demonstration that greater proximity to the behavioral consensus is associated with better objective performance. At the whole-brain level, neural typicality during the social movie was positively associated with CFMT accuracy (20,853 voxels after cluster correction; peak r = 0.60; N = 49; Supplementary Figure S4g) in regions including the bilateral fusiform gyri, superior and middle temporal gyri, cingulate cortex, inferior frontal gyrus, inferior and superior parietal cortex, and early visual cortex. To test, in a hypothesis-driven way, whether typicality within face-processing regions specifically tracks behavior on face-related tasks, we computed each participant’s mean neural typicality in the left and right FFA (individually defined) and the left and right amygdala, and correlated it with all behavioral measures (Spearman, one-tailed in the predicted direction; Supplementary Table S1). During the social movie, typicality in both left and right FFA correlated with CFMT accuracy (ρ = 0.26 and 0.27), as did typicality in the left amygdala (ρ = 0.35). More broadly, FFA typicality was related exclusively to tasks involving faces or human bodies: left FFA typicality also correlated with social gaze typicality during both the animal and social movies (ρ = −0.25 and −0.26), with animal gaze typicality during both movies (ρ = −0.35 and −0.29), and with biological motion prediction accuracy during both the social (ρ = 0.32) and the object movie (ρ = 0.35). No task without faces (object gaze, car memory, non-biological motion prediction or sound ratings) correlated with FFA typicality in any movie. Notably, this included emotion-rating typicality, a strongly social but purely auditory task, which was related to neither FFA nor amygdala typicality in any movie, dissociating face processing from social content more generally. A direct test confirmed this specificity: in the left FFA, correlations with face(/body) tasks were stronger than with non-face tasks (mean aligned ρ = 0.19 vs −0.04; permutation p < .001; exact task-label permutation p = .008, the most extreme of all 126 possible task assignments; Supplementary Table S2), most clearly during the social (p < .001) and animal (p = .005) movies and weaker during the object movie (p = .057), and the result held when biological motion prediction was classified as a non-face task (p = .006) or excluded (p < .001). The amygdala, in contrast, was less selective for faces: its typicality was related to social gaze (bilaterally, animal movie), animal gaze (left, animal movie), biological motion prediction (right, social movie) and face recognition (left, social movie), but also to object gaze during the animal movie (ρ = −0.24 and −0.27), a task without faces. Individual correlations were consistent in direction but modest, and none survived FDR correction across the 36 tests per movie (lowest q = .18). Together with the motion-prediction results, these findings show that neural typicality predicts performance on multiple tasks with objectively defined correct responses, and that within the FFA this relationship is specific to tasks that engage face processing.

### Convergent Neural Correlates of Behavioral Typicality

The preceding analyses showed that different behavioral measures were associated with partially distinct, content-appropriate neural systems. We finally asked whether, against this specificity, any regions recur across behavioral domains. We therefore combined the cluster-corrected maps of nine brain–behavior analyses and counted, for each voxel, the number of analyses with a significant relationship (see Methods). Most voxels in the overlap map were associated with two analyses (80,558 voxels), and the number of voxels decreased steeply with the number of overlapping analyses (three: 32,256; four: 11,105; five: 2,759; six: 680; seven: 25), with no voxel associated with more than seven of the nine analyses (Figure 6). The regions shared by the most measures were predominantly left-lateralized, including the posterior superior temporal sulcus and middle temporal gyrus, inferior frontal gyrus, supramarginal and angular gyri, dorsolateral prefrontal cortex, and the medial paracentral and supplementary motor region. Thus, alongside the content-specific relationships observed across tasks, a smaller set of regions was consistently associated with individual differences across multiple behavioral domains.

**Figure 6.**
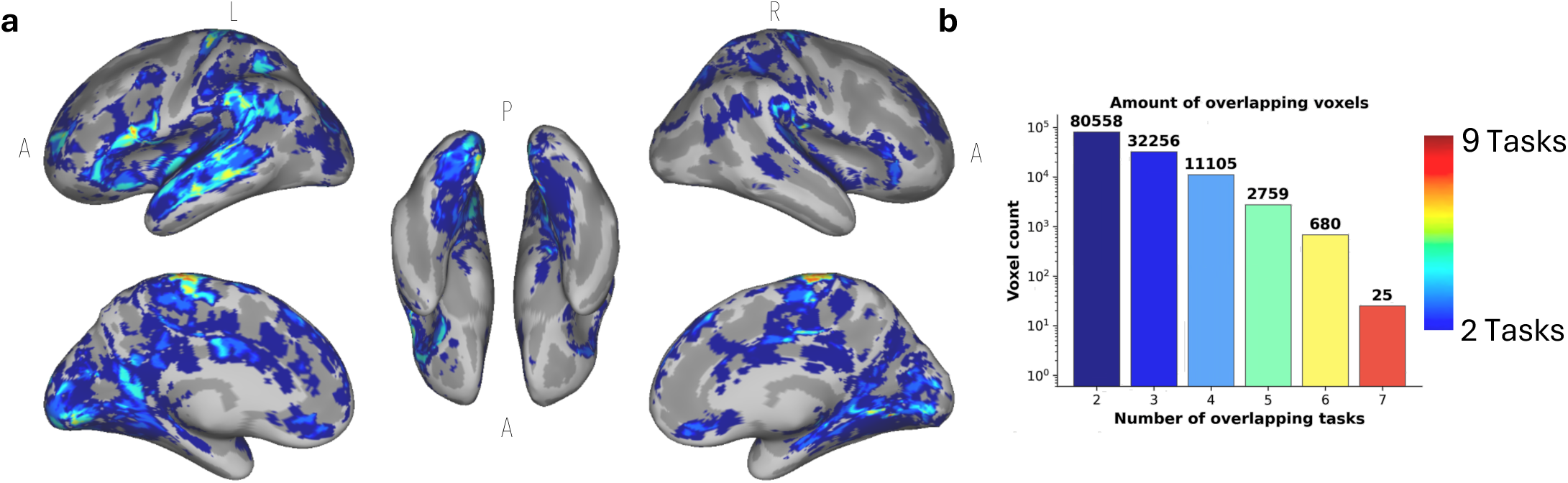
Convergent neural correlates across behavioral domains. **(a)** number of brain–behavior analyses with a significant relationship at each voxel, shown for voxels associated with at least two analyses. Nine brain–behavior maps were combined: social-movie neural typicality related to social, animal and object gaze typicality, emotion-rating typicality, biological and non-biological motion prediction accuracy, and CFMT and CCMT accuracy, and object-movie neural typicality related to non-biological motion prediction accuracy. Each map was cluster-corrected separately with its own cluster-extent thresholds (see Methods) and dilated by one voxel before summation to accommodate minor spatial misalignment; voxel counts in (b) are computed from the dilated maps. Warmer colors indicate voxels associated with more measures (2–7). **(b)** Number of voxels associated with exactly n measures (log scale).

## Discussion

Naturalistic stimuli reliably evoke shared responses across individuals, and much work has used this shared structure to study what people have in common and, more recently, how similarity between individuals relates to their behavior. Here we asked what an individual’s proximity to the shared response signifies. We found that typicality is a stable characteristic of the individual across both behavioral and neural measures, but that this stability is specific to the computation being performed, rather than reflecting a global tendency to respond like others. Moreover, in tasks with independently defined correct responses, proximity to the neural consensus during naturalistic viewing predicted performance. We argue that these findings are best understood if the consensus approximates a shared solution to the computation being performed, and consider how this view relates to work linking neural similarity between individuals to their relationships and behavior.

Reliable convergence across observers is well established for neural responses during naturalistic viewing (Hasson et al. 2004; Nastase et al. 2019), and reliability has been proposed as a core requirement for the measurement of individual differences (Kadlec et al. 2024). Our results show that individual positions relative to the consensus are similarly reliable. Behaviorally, typicality was highly reliable across independent stimuli within the same condition, but correlated significantly less across conditions engaging different processes, and was largely unrelated across tasks (Figure 2). Neural typicality showed a parallel organization: individual differences were preserved across movie categories, particularly within regions similarly engaged by the different stimuli, whereas regions preferentially engaged by a particular movie showed greater stimulus specificity (Figure 3; Supplementary Figure S3). This pattern argues against typicality reflecting a single global disposition, such as attentiveness, compliance, or a general tendency toward consensus, which would predict comparable typicality across tasks and conditions. Instead, an individual can be reliably typical within one context without being uniformly typical across all forms of processing.

### Computation Specificity Across Brain Systems and Stimuli

If typicality is defined relative to a particular computation, its relationship with behavior should also be specific, both in where it is expressed (within the systems supporting the computation) and in when (during stimuli that engage it). Because naturalistic movies simultaneously engage perceptual, attentional, social, emotional, mnemonic, and predictive processes (Hasson et al. 2010, 2004; Nastase et al. 2020), a single stimulus can reveal multiple stable characteristics of the same individual, each expressed within the systems supporting it (Finn et al. 2020; Ramot et al. 2020). Different behavioral measures were indeed associated with neural typicality in different, functionally relevant systems: social gaze typicality with regions supporting social orienting (Figure 4a; Supplementary Figure S7a), and emotion-rating typicality with regions involved in affective and social processing (Figure 4d; Supplementary Figure S7b). The face-patch ROI analysis made this specificity explicit (Supplementary Tables S1–S2). Typicality in the FFA was related to behavior only on tasks involving faces or bodies (face recognition, biological motion prediction, and gaze during social and animal movies), and a direct test confirmed that these relationships were stronger than those with non-face tasks, including the Cambridge Car Memory Test, despite its closely matched demands. This fits our previous finding that the network supporting individual differences in face memory did not predict memory for cars (Ramot et al. 2019). The amygdala, by contrast, was less face-selective: its typicality was also related to gaze during object movies. The pattern also dissociates faces from social content: gaze during animal movies, which contain faces but no human social interaction, was related to FFA typicality, whereas emotion ratings, a strongly social but purely auditory task, were not.

The same specificity was evident with respect to stimulus content, though less absolutely. Face-patch typicality was related to gaze and face-recognition performance only during the social and animal movies, although its relationship with biological motion prediction was also evident during the object movie. Emotion-rating typicality was related to neural typicality during the social movie, but showed fewer, directionally inconsistent relationships during the object movie. Gaze showed a graded dependence on content: social-movie neural typicality was most strongly related to gaze typicality during social viewing, less extensively during animal viewing, and only weakly during object viewing (Figure 4a–c), whereas object-movie neural typicality showed few, directionally inconsistent relationships with gaze typicality (Supplementary Figure S5). Finally, performance on both motion-prediction tasks was related to neural typicality in widespread regions during both movies; the extent of these relationships differed between movies, but a direct test of a task × movie crossover was not significant, indicating that the motion-prediction results do not by themselves establish stimulus specificity.

These results echo recent evidence that both intersubject correlation and its associations with behavior vary considerably across movies (Leipold et al. 2024). Such variability has primarily been framed as a challenge to the generalizability of naturalistic findings. Our results suggest that it may, at least in part, be expected: if the consensus reflects a shared solution to the computations a stimulus engages, proximity to it should be informative about behavior mainly when the stimulus engages the computation on which that behavior depends. Consistent with this, the stimulus specificity observed here for gaze, emotion ratings, and face-related behavior followed the correspondence between stimulus content and task demands. Stimulus specificity may therefore be not only a limitation of naturalistic paradigms, but also a tool, allowing stimuli to be selected to probe particular computations.

### What Does the Group Consensus Represent?

Shared responses during naturalistic viewing reflect the computations engaged by the stimulus, and track shared memories and interpretations (Hasson et al. 2004; Lahnakoski et al. 2012; Chen et al. 2017; Yeshurun et al. 2017; Nguyen et al. 2019). Our results suggest that an individual’s proximity to these shared responses may also carry information about how well those computations are performed. Efficient coding theories propose that neural representations are shaped to make the best use of limited resources given the statistics of the environment (Simoncelli and Olshausen 2001; Barlow 2012). Individuals exposed to similar environmental statistics, and operating under similar biological constraints and objectives, would then be expected to converge independently on similar solutions, while idiosyncratic deviations that are not systematically shared are averaged out in the consensus (Galton 1907; Surowiecki 2004). The consensus may thus approximate an effective, or even optimal, solution to the computation, with typicality indexing how closely an individual approaches it.

The tasks with objective performance measures allowed us to test this. In both motion prediction and face recognition, proximity to the behavioral consensus was strongly associated with accuracy. Because these were forced-choice tasks, this is expected to some degree: when most observers respond correctly, the consensus and the correct response largely coincide. More informative is that neural typicality, measured during passive movie viewing, predicted performance on these tasks, which were performed at a different time and in a different context.

These findings do not imply that the consensus is inherently optimal, or that deviation from it necessarily reflects poorer processing. Averaging preserves whatever is consistently shared, including shared biases, and naturalistic stimuli may permit multiple successful interpretations. The consensus should therefore approximate an optimal solution only to the extent that individuals share the same objectives and constraints. Where they do, typicality may provide an empirical approximation of successful processing in contexts where objective ground truth cannot be defined. Where they do not, the consensus itself may differ across groups, a possibility we consider next.

### Consensus, Similarity, and Group Structure

A growing body of work has related similarity between individuals’ neural responses to their relationships and characteristics. Friends respond more similarly than more distant acquaintances (Parkinson et al. 2018), as do sisters and spouses (Li et al. 2022; Bacha-Trams et al. 2024), and individuals with similar political views or personality traits (Matz et al. 2022; Leong et al. 2020; Van Baar et al. 2021). Inter-subject representational similarity analyses have extended this to performance, showing that high performers on cognitive tasks tend to be more synchronized with one another, whereas low performers are idiosyncratic in their own ways (Finn et al. 2020; Chamberlain et al. 2024). This “Anna Karenina” pattern is what would be expected if high performers lie close to a shared consensus, and is in this sense consistent with the present findings. Notably, it was observed for task-based measures but not for questionnaire-based measures (Chamberlain et al. 2024), in line with the idea that convergence is most informative for computations with a well-defined solution, although the two types of measures differ in many other respects. These analyses relate performance to how similar individuals are to one another. The present approach instead treats the group consensus itself as meaningful, approximating a shared solution to the computation at hand, and asks whether proximity to it predicts performance.

The relationship-based findings may at first appear to challenge the view of the consensus as an approximation of an optimal solution: if neural similarity tracks group membership, the consensus might simply reflect the average of whichever individuals happen to be sampled. We suggest instead that these findings are complementary to ours, and expected under the same framework. The assumption of shared objectives and constraints is unlikely to hold exactly. Individuals who share experiences, priors, or goals may effectively optimize for slightly different objectives than others, and would therefore be expected to converge on slightly different solutions. In efficient coding terms, systematic differences in experience would shift the statistics to which representations are adapted, and hence the solution toward which they converge. Individuals high in trait paranoia, for example, were more synchronized with one another during an ambiguous social narrative (Finn et al. 2018), consistent with a shared interpretive prior shaping a shared response. We would expect such group-level differences to be small relative to the response shared across all observers, shifting the consensus only slightly.

This account makes a prediction that the present study did not test: segmenting individuals according to their objectives, or the strategies they adopt, should yield more coherent subgroup consensuses, and possibly stronger relationships between typicality and performance. Such a structure would refine, rather than undermine, the interpretation of the consensus as a proxy for an effective solution. The consensus across a heterogeneous sample would then approximate an average of closely related solutions, and would remain informative to the extent that these solutions are similar.

### Convergence Across Computations

Although the brain–behavior relationships identified here were largely computation-specific, the underlying computations do not operate in isolation. A smaller set of regions, predominantly in the left hemisphere, recurred across multiple behavioral measures (Figure 6). These included the posterior superior temporal sulcus and middle temporal gyrus, the inferior frontal gyrus, the supramarginal and angular gyri, the dorsolateral prefrontal cortex, and the medial paracentral/supplementary motor region. Several of these regions overlap with areas previously described as cortical hubs, which participate flexibly across multiple functional networks and tasks (Van Den Heuvel and Sporns 2013). The present overlap analysis cannot establish such a role, but the recurrence of these regions across independent behavioral measures raises the possibility that they support processes shared across multiple aspects of naturalistic cognition.

### Typicality as a Framework for Studying Individual Differences

Typicality also offers a complementary approach to studying stable individual differences. Resting-state functional connectivity captures trait-like individual differences that generalize beyond any particular task (Finn et al. 2015; Tavor et al. 2016; Gratton et al. 2018). Neural typicality shows comparable stability across independent stimuli while remaining directly linked to ongoing computation, so that individual differences can be interpreted in the context of the processing being performed. Because naturalistic stimulation provides a common temporal structure across observers, typicality can be estimated at the voxel level without predefined regions or parcellations, allowing computation-specific signatures to emerge from the data, while remaining compatible with hypothesis-driven analyses such as the face-patch analysis presented here.

An important question is how well these relationships generalize. This is particularly relevant for typicality, where the reference itself is defined by convergence across the sampled observers. Our previous work provides some evidence that the reference can generalize across populations: autistic and typically developing participants converged on highly similar average gaze patterns, despite the autistic group lying further from this shared response (Ramot et al. 2020). Moreover, although the present imaging cohort is relatively modest, the relationship between social gaze typicality and neural typicality closely replicated our previous findings in an independent cohort, on a different scanner, and with different stimuli in a different language. The broader computation-specific relationships identified here will require confirmation in larger samples. In addition, each movie category was represented by a single movie in the present analyses, so differences between movies cannot be attributed to their content alone (Leipold et al. 2024); sampling multiple movies per category would allow content-specific effects to be separated from movie-specific ones. Moreover, although movies were distributed across sessions, the present design does not systematically separate within-from across-session comparisons; a formal test of the stability of neural typicality across sessions will be important for establishing it as a trait-like characteristic. Finally, it remains open how expertise shapes typicality. Expertise may bring responses closer to the shared solution, or, as a case of the subgroup structure discussed above, lead experts to converge on a different consensus as specialized objectives and strategies emerge. Clinical and neurodevelopmental conditions raise the complementary question. Altered biological constraints, whether developmental or imposed by neurological disease, may prevent individuals from reaching the shared solution, as suggested by the greater distance of the autistic participants described above from an otherwise shared gaze response, or may shift the solution itself. Because typicality is computation-specific and does not require task-specific ground truth, it may help identify which computations are affected in a given condition, and how they change over time.

Rather than treating shared responses and individual differences as separate descriptions of brain function (Vanderwal et al. 2019; Finn et al. 2020), the present findings suggest that they are two aspects of the same framework. The consensus provides the reference against which individual differences are quantified and, when observers share objectives and constraints, may approximate a shared solution to the computation being performed. Typicality quantifies how individuals diverge from it, in a manner specific to that computation. Similarity-based findings, including convergence within groups, fit within this view rather than competing with it. Together, these measures provide a principled way to study meaningful individual differences across behavioral and neural domains under naturalistic conditions, including where objective ground truth is unavailable.

## Methods

### Ethics

This study was carried out in accordance with the Weizmann Institutional Review Board (Weizmann IRB 3228-1) and Tel Aviv Souraski Medical Center Helsinki committee (TLV-0410-21) and complies with all relevant ethical regulations. All participants gave informed consent before starting the experiment.

### Participants

A total of 103 participants completed the behavioral battery (age 27±5; 37 females). Among them, 53 participants additionally underwent functional MRI (fMRI) scanning (age 27±5; 19 females). All participants had normal or corrected-to-normal vision and provided written informed consent prior to participation.

Participants completed the behavioral battery across three sessions, each lasting approximately 120 minutes. A subset of participants subsequently completed two fMRI scanning sessions comprising resting-state, movie-viewing, and task-based paradigms corresponding to the behavioral battery.

### Behavioral Battery

To measure social cognition across multiple domains, we designed a behavioral battery consisting of four tasks, including both well-established paradigms and newly developed tasks. Each task included both social and non-social stimulus conditions. The battery included Movie watching with eye tracking, emotion and sound intensity rating, motion prediction, and the Cambridge Face Memory Test (CFMT), the Cambridge Car Memory Test (CCMT). Stimuli were presented on a 60-Hz display.

Behavioral sessions were conducted in a quiet testing environment, and all data were processed using custom MATLAB scripts.

### Movie Watching with Eye Tracking

Participants viewed 180 short movie clips, each lasting 14 seconds, depicting three stimulus categories: human social interactions, behaving animals, and moving objects, with 60 clips per category. Movies were presented across three experimental sessions, with 60 clips per session and equal representation of each category. Stimuli were presented in randomized order with 6-second inter-trial intervals (see Figure 1a).

Eye movements were recorded using a Tobii Pro Spectrum eye tracker (Tobii, Danderyd, Sweden) at a sampling rate of 600 Hz. Participants were seated approximately 60 cm from the display, and Tobii’s built-in 5-point calibration was performed immediately prior to each experimental run (validating that average error was below 0.3 degrees).

### Gaze Preprocessing

Raw gaze coordinates were converted from normalized display coordinates into pixel space. Blink periods and high-velocity artifacts were identified and removed, and off-screen samples were excluded from further analyses.

### Gaze Typicality

To quantify individual differences in visual sampling during naturalistic viewing, we computed a leave-one-out gaze typicality measure following approaches used in prior work on inter-subject gaze alignment. For each participant and each movie, a reference gaze trajectory was computed as the mean gaze position of all other participants at each time point. The participant’s gaze at each time point was then compared to this reference by computing the Euclidean distance between them, and these distances were averaged across the duration of the movie.

This procedure yielded a movie-level typicality score reflecting the degree to which an individual’s gaze aligned with the group-consensus viewing pattern. Scores were subsequently averaged within stimulus categories to obtain category-specific gaze typicality measures. Higher values indicate greater deviation from the normative gaze trajectory. This approach captures the extent to which individuals sample visual information in a manner consistent with shared attentional priorities of the group during naturalistic viewing, and builds on prior work linking gaze typicality to social orienting mechanisms (Ramot et al. 2020).

To evaluate the robustness of the gaze typicality measures, we assessed the split-half reliability of each movie by repeatedly dividing participants into two independent groups, computing the group-average gaze pattern for each half, and correlating the resulting trajectories (Supplementary Figure S2a). This procedure was repeated 1,000 times. Movies with split-half reliability below r = 0.8 were excluded, removing 19 of the 180 movies. From the remaining 161 movies, we selected a subset matched across categories on the mean and variance of participants’ typicality values, yielding 133 movies (social = 45, object = 43, animal = 45; Supplementary Fig. S2). Only these 133 movies were used in all subsequent analyses.

### Gaze Typicality Clustering Analysis

Per-movie gaze typicality profiles across participants (45 animal, 45 social, 43 object movies) were k-means clustered (correlation distance; missing values imputed with each movie’s across-participant mean). Silhouette analysis (k = 2–10) supported k = 3, matching the three stimulus domains. Cluster purity — the proportion of movies in a domain assigned to that domain’s modal cluster — was computed per domain; object-segment purity was tested against a null distribution from 10,000 random 45-movie samples (Supplementary Figure S1).

To test whether animal movies reliably grouped with the social-typicality cluster, clustering was repeated 1,000 times; animal-movie cluster labels were used to build a co-clustering consensus matrix, which was hierarchically clustered (average linkage) to yield a stable animal-movie cluster assignment, compared against the social cluster from the primary solution.

### Emotions Intensity Evaluation Task

Participants listened to 18 audio clips containing either conversational speech or environmental sounds (1-2 minutes each). After each clip, participants rated short ( < 4 seconds) segments, taken from the full clip, on a 1–7 Likert scale. For speech stimuli, ratings reflected perceived emotional intensity of a given emotion, whereas for environmental sounds, ratings reflected the intensity of specific acoustic features (see Figure 1b). Participants rated 218 emotions and 226 environmental sounds overall. This task was designed to capture individual differences in sensitivity to socially relevant auditory information, drawing on prior work demonstrating that vocal cues contribute to empathic understanding and shared physiological responses (Kraus 2017; Jospe et al. 2020). Behavioral typicality was computed using a leave-one-out approach in which each participant’s ratings were compared to the group-average ratings of all other participants, and the absolute deviation was averaged across stimuli.

To evaluate the robustness of these ratings, we assessed split-half reliability by repeatedly dividing participants into two independent groups, computing the group-average rating for each half, and correlating the resulting scores across repeated random splits (Supplementary Fig. S2). Consensus ratings showed high split-half reliability for both emotional speech (mean r = 0.97, SD = 0.004, 95% CI [0.96, 0.98]) and environmental sounds (mean r = 0.99, SD = 0.002, 95% CI [0.98, 0.99]), and no clips or segments were excluded from subsequent analyses.

### Movement Prediction Task

Participants viewed short video clips depicting motion either by human agents about to perform a hand movement to the left/right or about to jump to the left/right (biological motion) or by inanimate objects (rocks/leaves) about to be moved by wind to the left/right (non-biological motion; see Figure 1c). The clips were truncated to include only the initial frames preceding the onset of motion, thereby providing minimal cues regarding the upcoming direction of movement. Participants were required to intuitively predict the direction of motion, indicating whether it would proceed to the left or right. This design builds on prior work showing that observers can extract predictive information about actions and future states from limited visual input and partial scene information (Hart et al. 2020; Vaziri-Pashkam et al. 2017).

To quantify prediction typicality, we computed, for each trial, the proportion of participants who selected each response option (left/right). Each participant’s typicality score for that trial was defined as the proportion of participants whose responses differed from their own, such that responses aligned with the majority yielded low deviation values, whereas responses aligned with the minority yielded high deviation values. Trial-level deviation values were averaged across trials within each domain to yield participant-level typicality scores separately for biological and non-biological motion trials. Prediction accuracy was computed separately by comparing each participant’s response to the objectively correct direction of motion on each trial.

To evaluate the robustness of these measures, we assessed reliability at two levels. First, to confirm that item difficulty was itself a stable property of the trials rather than an artifact of who was tested, we repeatedly split participants into two independent groups, computed each group’s average accuracy for every trial, and correlated the two groups’ trial-by-trial accuracy patterns. Second, to assess whether individual participants’ accuracy and typicality scores were themselves stable, we repeatedly split trials (rather than participants) into two independent subsets, computed each participant’s accuracy or typicality score within each subset, and correlated the resulting scores across participants, separately within each domain. Both measures showed high split-half reliability for biological motion (accuracy: mean r = 0.82, SD = 0.03, 95% CI [0.75, 0.87]; typicality: mean r = 0.83, SD = 0.03, 95% CI [0.77, 0.88]) and non-biological motion (accuracy: mean r = 0.91, SD = 0.02, 95% CI [0.88, 0.94]; typicality: mean r = 0.91, SD = 0.02, 95% CI [0.88, 0.94]), and no trials or clips were excluded from subsequent analyses. Item difficulty was similarly stable across independent groups of participants for both biological (mean r = 0.91, SD = 0.01, 95% CI [0.89, 0.93]) and non-biological motion (mean r = 0.85, SD = 0.02, 95% CI [0.81, 0.88]), with no relationship between domains (mean r = 0.02, SD = 0.02, 95% CI [−0.03, 0.06]) (Supplementary Figure S2).

### Cambridge Memory Tests

Participants completed the Cambridge Face Memory Test and the Cambridge Car Memory Test, which assess recognition memory for faces and objects, respectively. Performance was quantified as the total number of correct responses across trials, following standard scoring procedures (Duchaine and Nakayama 2006; Dennett et al. 2012).

To quantify behavioral typicality for the CFMT, we additionally computed, for each trial and subject, the proportion of the full sample that selected the same response as that subject; typicality was defined as one minus this proportion, such that higher scores reflect more atypical (less commonly given) responses, and per-subject typicality was averaged across all trials.

### Behavioral Statistical Analysis

For all tasks, individual differences were quantified as typicality, each participant’s deviation from the group consensus. For continuous measures (gaze, emotion and sound ratings), the consensus was computed from all other participants (leave-one-out). For the forced-choice tasks (motion prediction, CFMT), typicality was defined from the response distribution of the full sample, in which each participant’s own response contributes less than 1% of each response proportion. Reliability of individual scores was assessed by split-half analysis across stimuli: stimuli within each category were randomly divided into two halves, scores were computed separately for each half, and the two sets of scores were correlated across participants. This was repeated over 10,000 random splits. Chance level was estimated by repeating the procedure after randomly permuting participant identities between halves, taking the 95th percentile of the resulting null distribution. To test category specificity, for each random split we computed the difference between the within-category split-half correlation and the corresponding cross-category correlation (scores from one half of category A correlated with scores from one half of category B). We report mean differences, 95% intervals (2.5th–97.5th percentiles across the 10,000 splits) and two-sided p-values (the proportion of splits in which the difference fell on the opposite side of zero). The difference in the size of the within-versus between-category gap between social and non-social categories was tested in the same way. Finally, to test whether typicality reflected a general disposition across tasks, we computed Pearson correlations between all pairs of the nine behavioral measures from different tasks (30 comparisons), with Benjamini–Hochberg FDR correction (Benjamini and Hochberg 1995).

### MRI Methods

#### MRI Data Acquisition

A subset of 53 participants from the behavioral cohort completed two functional magnetic resonance imaging (fMRI) sessions. Data were acquired using a 7 Tesla Siemens Terra MRI scanner equipped with a 32-channel head coil, at the neuroimaging center in the Weizmann Institute of Science. Functional MR images were acquired with a multiband echo planar imaging and multi-echo gradient sequence (TR = 2.01 s; flip angle 70°; echo times 13.2, 34.72, and 56.2 ms; number of slices 72; voxel size 1.6 x 1.6 x 1.6 mm; multi-band acceleration factor = 3; GRAPPA = 3; Bandwidth = 2056 Hz/Px). T1-weighted anatomical images were acquired using a 3D MP2RAGE sequence (TR = 4.46 s; voxel size 1 x 1 x 1 mm; flip angle = 4°; TE = 2.19 ms; GRAPPA = 3; Bandwidth = 200 Hz/Px).

#### fMRI Paradigm

Each of the two scanning sessions began with two 8-min resting-state runs, followed by movie viewing and task-based paradigms. Naturalistic movie viewing consisted of 5-minute movie segments from three categories – social/animals/objects (see Figure 1a). The movies were randomly distributed between the sessions. Task-based paradigms were selected to parallel the behavioral battery and included emotion intensity rating (2 trials in each condition; 16 minutes) and movement prediction (140 trials; biological/non-biological motion; 12 minutes). Structural anatomical images were acquired in each session for registration and normalization procedures.

Participants were instructed to remain still throughout scanning and were monitored continuously for compliance, awareness, and alertness.

#### fMRI Preprocessing

Preprocessing was performed using AFNI (Cox 1996). Functional datasets underwent standard preprocessing procedures including removal of first 4 TRs, slice-timing correction, motion correction, spatial normalization, and transformation into MNI space using AFNI’s sswarper (Saad et al. 2009), after denoising the MP2RAGE image with MPRAGEise.py (S. Kashyap; https://github.com/srikash/MPRAGEise). Multi echo data was preprocessed using ME-ICA/tedana (v25.0; Kundu et al. 2012; DuPre et al. 2021; Kundu et al. 2013).

To reduce high-frequency spatial noise while preserving anatomical specificity, spatial smoothing was applied to the movie-viewing data, using a 4-mm full-width-at-half-maximum (FWHM) Gaussian kernel restricted to brain voxels. Preprocessed residual time-series datasets were stored in AFNI BRIK/HEAD format and used for all subsequent analyses.

For further typicality assessments, to ensure comparable spatial coverage across participants, a group mask was generated by retaining voxels containing valid signal in at least 80% of participants in each movie. All voxelwise analyses were restricted to this common mask.

#### Quality Control and Participant Exclusion

Several quality-control procedures were implemented to ensure data quality and participant compliance.

Head motion was assessed for all scanning runs, and motion estimates were included as nuisance regressors in all first-level analyses. No participant was excluded for head motion (mean framewise displacement < 0.2 mm for all participants). Participant engagement was assessed from the neural data. For each movie, we identified voxels showing reliable shared responses across the full sample (group-mean neural typicality, |r| > 0.17) and computed each participant’s mean neural typicality within these voxels. Participants whose mean typicality fell more than 2 SD below the sample mean were considered not to have engaged with the movie and were excluded from that movie’s neural analyses (social movie: z = −2.41 and −2.37; object movie: z = −2.50 and −2.27). A third participant in the object movie fell just beyond this cutoff (z = −2.22) and was retained. For the animal movie, two participants had no usable data. This yielded N = 51 per movie.

#### Neural Typicality During Movie Viewing

Neural typicality was quantified using a leave-one-out inter-subject correlation (ISC) approach.

For each participant, voxelwise BOLD time series recorded during the 5-min movie viewing were extracted and compared with the average time series of all remaining participants. Specifically, for each voxel, the participant’s time series was correlated with the leave-one-out group-average time series, yielding a whole-brain map of neural typicality. Higher values indicated greater similarity between an individual’s neural responses and the group-consensus response, whereas lower values reflected more idiosyncratic neural processing. Pearson correlations were used (N = 51 per movie).

Neural typicality maps were computed separately for each movie, allowing examination of both stimulus-specific and domain-general individual differences in neural processing.

Statistical significance was estimated separately for each movie using a time-reversal null model, yielding null-derived correlation thresholds of |r| = 0.076 for the social movie, |r| = 0.064 for the object movie, and |r| = 0.099 for the animal movie.

#### Neural Stability Analyses

To assess the reliability of neural typicality measures within and across domains, several complementary stability analyses were performed.

#### Within-Movie Stability

To assess whether individual neural typicality was stable within a movie, each participant’s movie time series was divided into two independent halves (the first 75 and the last 76 TRs), and neural typicality was computed separately for each half, using a leave-one-out reference computed within that half. For each voxel, the two halves’ typicality values were then correlated across participants (Pearson; N = 51 per movie), yielding a map of within-movie stability. Significance was assessed by permutation (5,000 permutations), in which the pairing of participants between the two halves was randomly shuffled and the stability map recomputed. Voxelwise maps were thresholded at p < .05, .01, .005 and .001, and cluster-size thresholds were derived from the pooled null distribution of cluster sizes (clusters of at least 6 voxels; NN = 1), taking the 99th percentile for each voxelwise threshold. As in the main analyses, the four cluster-corrected maps were combined into a single map. Analyses were restricted to each movie’s group mask.

#### Between-Movie Stability

To assess whether individual differences in neural typicality generalized across stimulus domains, typicality maps from each pair of movies were correlated voxelwise across participants with typicality data for both movies (N = 47–49 per pair), and significance was assessed with the same permutation and cluster-correction procedure (5,000 permutations of the participant pairing between movies; 99th percentile; clusters of at least 6 voxels). Analyses were restricted to a common mask across movies.

#### Brain–Behavior Correlation Analyses

To identify neural correlates of behavioral individual differences, voxelwise correlations were computed between neural typicality maps and behavioral measures. For each behavioral measure, outliers were identified within the fMRI sample (N = 53): participants more than 2 SD from the mean were excluded for typicality measures, and participants more than 2 SD below the mean for accuracy measures (high performers were retained). Participants excluded from one motion-prediction task were excluded from both. For the remaining participants, subject-specific neural typicality maps were concatenated across participants and correlated with the corresponding behavioral scores using AFNI’s 3dTcorr1D. This analysis generated whole-brain statistical maps highlighting regions in which neural typicality was correlated with behavioral performance.

Brain-behavior analyses were conducted separately for each behavioral measure and movie condition. Both social and non-social behavioral metrics were examined to determine whether neural typicality exhibited domain-specific or domain-general relationships with behavior.

#### Statistical Inference

Brain–behavior correlation maps were assessed with a nonparametric, permutation-based cluster-extent correction. For each analysis, behavioral scores were randomly permuted across participants 5,000 times, and the voxelwise correlation map between neural typicality and the permuted scores was recomputed for each permutation, generating a null distribution under the hypothesis of no brain–behavior relationship. Each permuted map was thresholded at four voxelwise significance levels (two-tailed p < .05, .01, .005 and .001; correlation thresholds derived from the t distribution with N − 2 degrees of freedom), separately for positive and negative correlations, and the sizes of all resulting clusters were recorded (face-connected neighbors, NN = 1). For each voxelwise threshold, cluster sizes were pooled across permutations, clusters smaller than 20 voxels were discarded, and the 95th percentile of the remaining distribution was taken as the cluster-extent threshold. The observed correlation map was thresholded in the same way with AFNI’s 3dClusterize, and a voxel was considered significant if it belonged to a surviving cluster at any of the four voxelwise thresholds; the four cluster-corrected maps were combined into a single map. Across all brain–behavior analyses, cluster-extent thresholds ranged from 130–165 voxels (p < .001), 167–199 voxels (p < .005), 193–220 voxels (p < .01) and 287–310 voxels (p < .05). For the partial correlation analysis, permutations followed the Freedman–Lane procedure and voxelwise thresholds used N − 3 degrees of freedom (see Partial Correlation). The within- and between-movie stability analyses used the same framework, permuting the pairing of participants between halves or movies, with a minimum cluster size of 6 voxels and a 99th-percentile cluster-extent threshold. Group-average typicality maps were thresholded separately using the time-reversal null described below. All reported results survived the relevant correction unless otherwise stated.

To assess robustness, we additionally applied a conservative family-wise criterion, using a cluster-forming threshold of p < .001 and a cluster-extent threshold taken from the 95th percentile of the maximum null cluster size across permutations. Under this criterion, the relationships of social-movie neural typicality with social gaze typicality and with biological motion prediction accuracy, and of object-movie neural typicality with non-biological motion prediction accuracy, remained significant (564, 566 and 371 voxels, respectively).

#### Overlap Analyses

To identify regions associated with multiple brain–behavior relationships, we combined the cluster-corrected binary masks (95th-percentile permutation threshold; see Statistical Inference) of nine brain–behavior analyses: social-movie neural typicality related to social, animal and object gaze typicality, emotion-rating typicality, biological and non-biological motion prediction accuracy, and CFMT and CCMT accuracy, and object-movie neural typicality related to non-biological motion prediction accuracy. Non-biological motion prediction thus contributes two maps, which reflect its relationships with neural typicality during two different movie categories. Each mask was dilated by one voxel (AFNI 3dmask_tool, 18-neighbour connectivity) to accommodate minor spatial misalignment across analyses, and the dilated masks were summed to give, for each voxel, the number of analyses (0–9) showing a significant brain–behavior relationship. Because each input mask was already cluster-corrected, no further threshold was applied to the summed map. Voxels associated with at least two analyses are displayed, and voxel counts for each level of overlap were computed from this same dilated map; these counts therefore include voxels within one voxel of a significant cluster.

#### Group-Average Typicality Maps and Stimulus-Domain Overlap

For each movie, a null distribution of typicality was estimated by time-reversing each participant’s time series and correlating it against the (non-reversed) leave-one-out group-average time series, yielding a null typicality map that preserved the temporal autocorrelation of the data while eliminating genuine stimulus-locked inter-subject correspondence. Null typicality maps were Fisher z-transformed, averaged across participants, and back-transformed to r, producing one group-level null mean-r map per movie. The 95th percentile of |r| within each movie’s brain mask was taken as that movie’s null-derived voxelwise threshold.

Group-level typicality maps were computed identically from the true (non-reversed) per-participant typicality maps and thresholded using each movie’s own null-derived cutoff, yielding a group mean-r typicality map per movie in which effect size was preserved (rather than binarized) at all suprathreshold voxels.

For visualization of overlap across stimulus domains only, the top 70% of |r| values within each movie’s null-thresholded map were additionally selected as a descriptive subset of the strongest effects (not an independent statistical threshold). These top-70% masks were combined across the three movies (social, animal, and object) into a single overlap map, coded such that each voxel’s value indicated the specific combination of stimulus domains in which it survived.

#### Task × Movie interaction

To test whether motion-prediction accuracy was more strongly related to typicality during the movie whose content matched the task, we computed, within a common mask, voxelwise Fisher-z correlations between social-and object-movie typicality and biological and non-biological motion-prediction accuracy (N = 46), and averaged the crossover contrast [z(social, biological) − z(object, biological)] − [z(social, non-biological) − z(object, non-biological)] across voxels. Significance was assessed by randomly exchanging each participant’s social- and object-movie maps (10,000 permutations), which preserves each participant’s overall typicality. The test was repeated using mean typicality within independent motion-task activation masks. The corresponding simple effects (z(social, biological) − z(object, biological), and z(object, non-biological) − z(social, non-biological)) were tested in the same way.

#### Region-of-interest analysis

To test whether typicality in face-selective and limbic regions relates to behavior, we computed each participant’s mean neural typicality within the left and right FFA and the left and right amygdala, separately for each movie. FFA ROIs were defined individually as 6-mm spheres centered on the center of mass of each participant’s face-selective fusiform cluster (faces > objects localizer). Amygdala ROIs were bilateral anatomical masks defined from the Eickhoff–Zilles macro-label atlas in MNI space (Eickhoff et al. 2005). Means were computed over voxels with valid signal. ROI typicality was correlated with each behavioral measure using Spearman rank correlations (N = 46–49 after the exclusions described in Brain–Behavior Correlation Analyses), tested one-tailed in the predicted direction: negative for typicality measures (distances from the consensus) and positive for accuracy measures. FDR correction (Benjamini–Hochberg) was applied across the 36 tests (9 behavioral measures × 4 ROIs) per movie.

#### Test of face specificity

To test directly whether ROI typicality was more strongly related to face-related than to non-face behavior, we computed, for each ROI and movie, the difference between the mean correlation across face(/body) tasks (social and animal gaze typicality, biological motion prediction, CFMT) and the mean across non-face tasks (object gaze typicality, CCMT, non-biological motion prediction, emotion- and sound-rating typicality), after aligning signs so that positive values indicated the predicted direction. Significance was assessed by permuting participants’ ROI values relative to all behavioral measures jointly (10,000 permutations), preserving the correlations among behavioral measures, and was confirmed with an exact permutation of task labels (all 126 assignments of four ‘face’ labels among the nine tasks). Tests were one-sided (face > non-face), performed for each movie and for the average across movies.

#### Partial correlation

To test whether the relationship between social gaze typicality and neural typicality during the social movie was specific to social content, we computed the voxelwise partial correlation between neural typicality and social gaze typicality, controlling for animal gaze typicality (N = 48). Cluster-extent thresholds were derived from a dedicated permutation null (5,000 permutations, Freedman–Lane procedure, Freedman and Lane 1983)zvu: the residuals of social gaze typicality after regression on animal gaze typicality were permuted, and the partial correlation map was recomputed for each permutation. Voxelwise thresholds were based on partial-correlation degrees of freedom (N − 3), with the same percentile and minimum-cluster criteria as the main analyses (Supplementary Figure S4c).

#### Task-based GLMs

Task data (emotion rating and motion prediction) were preprocessed as described above, and subject-level responses were estimated with standard general linear models in AFNI (3dDeconvolve; duration-modulated block regressors for each task condition, with polynomial drift and motion nuisance regressors, and censoring of high-motion timepoints). Group maps for each condition against rest were obtained with one-sample t-tests (3dttest++; emotion N = 51, motion N = 52) and are shown in Supplementary Figure S6. For the overlap analyses, the group maps were thresholded at t > 3.29: the social-rating periods of the emotion task (Figure 4d) and biological and non-biological motion trials (Figure 5a–c). They were then intersected on the cortical surface with the corresponding brain–behavior maps.

#### Semantic decoding and network correspondence

To characterize the functional profile of each brain–behavior map, cluster-corrected maps (values outside significant clusters set to zero) were decoded against the Neurosynth database (version 7, term maps derived from study abstracts; Yarkoni et al. 2011) using NiMARE’s correlation decoder (Salo et al. 2023). For each of 93 curated cognitive terms, a meta-analytic association map was computed (MKDA chi-square; terms occurring in at least 0.1% of abstracts), and term relevance was quantified as the spatial correlation between that map and the brain–behavior map. Maps of typicality measures, which are distances from the consensus, were sign-inverted before decoding so that positive values corresponded to more typical behavior. The 20 terms with the highest correlations are shown for each map. Network correspondence was assessed with the Network Correspondence Toolbox (Kong et al. 2025) as the Dice overlap between each map’s significant voxels, resampled to the 2-mm MNI152 grid, and the networks of the Gordon et al. (2017) 17-network atlas.

#### Software

Behavioral preprocessing and analyses were performed with custom MATLAB scripts (MathWorks, Natick, MA, USA). fMRI preprocessing and analyses were performed with AFNI (v25.3.04; Cox 1996) and tedana (v25.0; DuPre et al. 2021), together with custom Python scripts (Python 3.13) using NumPy (2.1), SciPy (1.14) and NiBabel (5.2) for neural typicality, permutation testing, cluster correction and region-of-interest analyses. Volumetric results were projected onto standard-mesh cortical surfaces (std.60) using Connectome Workbench (v2.2.0; trilinear volume-to-surface mapping), which was also used to compute surface overlap maps and render brain figures. Plots were generated with Matplotlib (3.10) and MATLAB.

## Acknowledgements

We would like to thank all participants who took part in this study. We thank Dr. Edna Furman-Haran from the Weizmann Human Brain Imaging Center for optimizing the multi-echo fMRI sequence, and Natalie Oshri and Eiska Tegareh for help in fMRI data collection. We thank Micaela Feigelsohn, Liraz Hemo, and Lucia Pinkus for participant recruitment and help in data collection. We thank Maya Salomon-Hazut for helping with the fMRI preprocessing protocol. We thank Sasha Devore for insightful comments during the writing process. This research was generously supported by the European Research Council (ERC-2022-StG 101077921), Israel Science Foundation grant 829/22, the Irene and Jared M. Drescher Center for Research on Mental and Emotional Health, Knell Family Institute for Artificial Intelligence, the Mike and Valeria Rosenbloom Center for Research on Positive Neuroscience, the Zuckerman Center for Research on Learning, Memory, and Cognition, and the Zuckerman STEM leadership program. M.R. is the incumbent of the Roel C. Buck Career Development Chair, and a research associate program grant from the Council for Higher Education to M.W. The funders had no role in study design, data collection and analysis, decision to publish, or preparation of the manuscript.

## Data Availability

The participant-level neuroimaging and behavioral datasets generated and analyzed in this study are not publicly available because of restrictions related to participant privacy and informed consent. Source data underlying the figures and tables are available from the corresponding author upon reasonable request, subject to applicable ethical and data-sharing restrictions.

## Author contributions

M.Z., M.R., conceived and designed the study. M.Z., I.H., A.J., and M.W. collected the data. M.Z. performed the behavioral and neuroimaging analyses. M.Z., M.W., and M.R. interpreted the results and wrote the manuscript. All authors contributed to manuscript revision and approved the final version.

## Supplemantry

**Supplementary Table S1.** Correlations between mean neural typicality in face-selective and limbic regions and behavioral measures, for each movie.

| Measure / Region | Animal movie |  | Social movie |  | Object movie |  |
| --- | --- | --- | --- | --- | --- | --- |
| | $\rho$ | $p$ | $\rho$ | $p$ | $\rho$ | $p$ |
| **Social gaze typicality (predicted -)** | $N = 49$ | | $N = 49$ | | $N = 49$ | |
| Left FFA | **-0.25** | **.042** | **-0.26** | **.035** | 0.00 | .504 |
| Right FFA | -0.04 | .397 | -0.23 | .056 | 0.07 | .678 |
| Left amygdala | **-0.29** | **.022** | -0.23 | .053 | 0.22 | .935 |
| Right amygdala | **-0.31** | **.015** | -0.14 | .169 | 0.10 | .764 |
| **Animal gaze typicality (predicted -)** | $N = 49$ | | $N = 49$ | | $N = 50$ | |
| Left FFA | **-0.35** | **.008** | **-0.29** | **.020** | 0.03 | .570 |
| Right FFA | 0.00 | .491 | -0.10 | .257 | 0.22 | .934 |
| Left amygdala | **-0.26** | **.035** | -0.02 | .441 | 0.16 | .863 |
| Right amygdala | -0.20 | .081 | 0.10 | .743 | -0.04 | .401 |
| **Object gaze typicality (predicted -)** | $N = 49$ | | $N = 49$ | | $N = 50$ | |
| Left FFA | -0.14 | .161 | -0.16 | .140 | 0.08 | .716 |
| Right FFA | 0.02 | .566 | -0.04 | .404 | 0.23 | .944 |
| Left amygdala | **-0.24** | **.047** | 0.18 | .893 | 0.28 | .977 |
| Right amygdala | **-0.27** | **.032** | 0.23 | .940 | 0.21 | .933 |
| **Emotion-rating typicality (predicted -)** | $N = 48$ | | $N = 48$ | | $N = 48$ | |
| Left FFA | 0.10 | .743 | 0.07 | .671 | 0.11 | .767 |
| Right FFA | 0.13 | .812 | 0.06 | .666 | 0.00 | .498 |
| Left amygdala | 0.17 | .883 | -0.19 | .101 | 0.13 | .812 |
| Right amygdala | 0.31 | .983 | 0.01 | .525 | 0.27 | .968 |
| **Sound-rating typicality (predicted -)** | $N = 47$ | | $N = 47$ | | $N = 47$ | |
| Left FFA | 0.23 | .942 | 0.15 | .837 | 0.12 | .790 |
| Right FFA | 0.22 | .928 | 0.18 | .883 | 0.11 | .761 |
| Left amygdala | -0.08 | .289 | 0.02 | .549 | 0.05 | .620 |
| Right amygdala | 0.03 | .585 | 0.08 | .699 | 0.05 | .627 |
| **Biological motion prediction accuracy (predicted +)** | $N = 48$ | | $N = 48$ | | $N = 48$ | |
| Left FFA | 0.06 | .338 | **0.32** | **.014** | **0.35** | **.007** |
| Right FFA | -0.11 | .767 | 0.12 | .199 | 0.00 | .501 |
| Left amygdala | 0.05 | .362 | 0.23 | .054 | 0.06 | .350 |
| Right amygdala | -0.07 | .677 | **0.30** | **.019** | -0.09 | .732 |
| **Non-biological motion prediction accuracy (predicted +)** | $N = 48$ | | $N = 48$ | | $N = 48$ | |
| Left FFA | -0.03 | .568 | 0.03 | .420 | 0.12 | .199 |
| Right FFA | 0.01 | .464 | 0.04 | .398 | 0.07 | .308 |
| Left amygdala | -0.03 | .586 | 0.13 | .183 | 0.05 | .359 |
| Right amygdala | -0.16 | .861 | 0.16 | .145 | -0.23 | .944 |
| **Face memory accuracy (CFMT) (predicted +)** | $N = 48$ | | $N = 49$ | | $N = 48$ | |
| Left FFA | 0.11 | .233 | **0.26** | **.034** | 0.01 | .469 |
| Right FFA | 0.09 | .279 | **0.27** | **.031** | 0.21 | .074 |
| Left amygdala | 0.05 | .365 | **0.35** | **.007** | -0.05 | .631 |
| Right amygdala | -0.03 | .591 | 0.05 | .365 | -0.12 | .800 |
| **Car memory accuracy (CCMT) (predicted +)** | $N = 48$ | | $N = 48$ | | $N = 48$ | |
| Left FFA | -0.08 | .703 | -0.01 | .525 | -0.14 | .832 |
| Right FFA | -0.07 | .685 | 0.01 | .460 | 0.01 | .467 |
| Left amygdala | 0.14 | .171 | 0.06 | .333 | -0.12 | .793 |
| Right amygdala | -0.04 | .612 | -0.06 | .649 | -0.11 | .780 |

Note. Spearman rank correlations between each participant's mean neural typicality within the region of interest and their behavioral score, computed separately for each movie. FFA ROIs were individually defined 6-mm spheres centered on the center of mass of each participant's face-selective fusiform cluster (faces > objects localizer); amygdala ROIs were the left and right amygdala of the Eickhoff–Zilles macro-label atlas in MNI space. Means were computed over voxels with valid signal. Typicality measures are distances from the group consensus, so more typical behavior corresponds to lower values and the predicted correlation with neural typicality is negative (-); for accuracy measures the predicted correlation is positive (+). p-values are one-tailed in the predicted direction and uncorrected; values with $p < .05$ are shown in bold. None survived Benjamini–Hochberg FDR correction across the 36 tests per movie (lowest $q = 0.18$ ). N varies across measures because of outlier exclusion (see Methods). FFA, fusiform face area; CFMT, Cambridge Face Memory Test; CCMT, Cambridge Car Memory Test.

**Supplementary Table S2.** Test of face specificity: mean correlation for face(/body) versus non-face tasks in each region of interest.

| Region | Movie | Face tasks | Non-face tasks | Difference | p (perm.) | p (labels) |
| --- | --- | --- | --- | --- | --- | --- |
| **Left FFA** | Animal | 0.19 | -0.06 | **0.25** | **.005** | .024 |
|  | Social | 0.28 | -0.01 | **0.29** | **< .001** | .008 |
|  | Object | 0.08 | -0.07 | 0.15 | .057 | .103 |
|  | *All movies* | 0.19 | -0.04 | **0.23** | **< .001** | .008 |
| **Right FFA** | Animal | 0.01 | -0.09 | 0.09 | .174 | .087 |
|  | Social | 0.18 | -0.03 | **0.21** | **.010** | .008 |
|  | Object | -0.02 | -0.05 | 0.03 | .381 | .381 |
|  | *All movies* | 0.06 | -0.06 | **0.11** | **.022** | .056 |
| **Left amygdala** | Animal | 0.16 | 0.05 | 0.11 | .122 | .159 |
|  | Social | 0.21 | 0.04 | **0.17** | **.030** | .048 |
|  | Object | -0.09 | -0.11 | 0.01 | .444 | .444 |
|  | *All movies* | 0.09 | -0.01 | **0.10** | **.034** | .008 |
| **Right amygdala** | Animal | 0.10 | -0.05 | 0.16 | .052 | .135 |
|  | Social | 0.10 | -0.04 | 0.14 | .065 | .103 |
|  | Object | -0.07 | -0.18 | 0.10 | .129 | .064 |
|  | *All movies* | 0.04 | -0.09 | **0.13** | **.007** | .008 |

*Note.* Values are mean direction-aligned Spearman correlations (signs flipped for typicality measures so that positive values indicate the predicted direction) across face(/body) tasks (social and animal gaze typicality, biological motion prediction, CFMT) and non-face tasks (object gaze typicality, CCMT, non-biological motion prediction, emotion- and sound-rating typicality), and their difference. p (perm.): one-sided permutation test shuffling participants' ROI values relative to all behavioral measures jointly (10,000 permutations). p (labels): exact permutation of task labels (126 possible assignments; minimum attainable $p = .008$ ). 'All movies' averages the three movies. Differences with $p$ (perm.) $< .05$ are shown in bold. Robustness, left FFA, all movies: with biological motion prediction classified as a non-face task, difference = 0.16, $p = .006$ ; with biological motion prediction excluded, difference = 0.21, $p < .001$ .

**Supplementary Figure S1.**
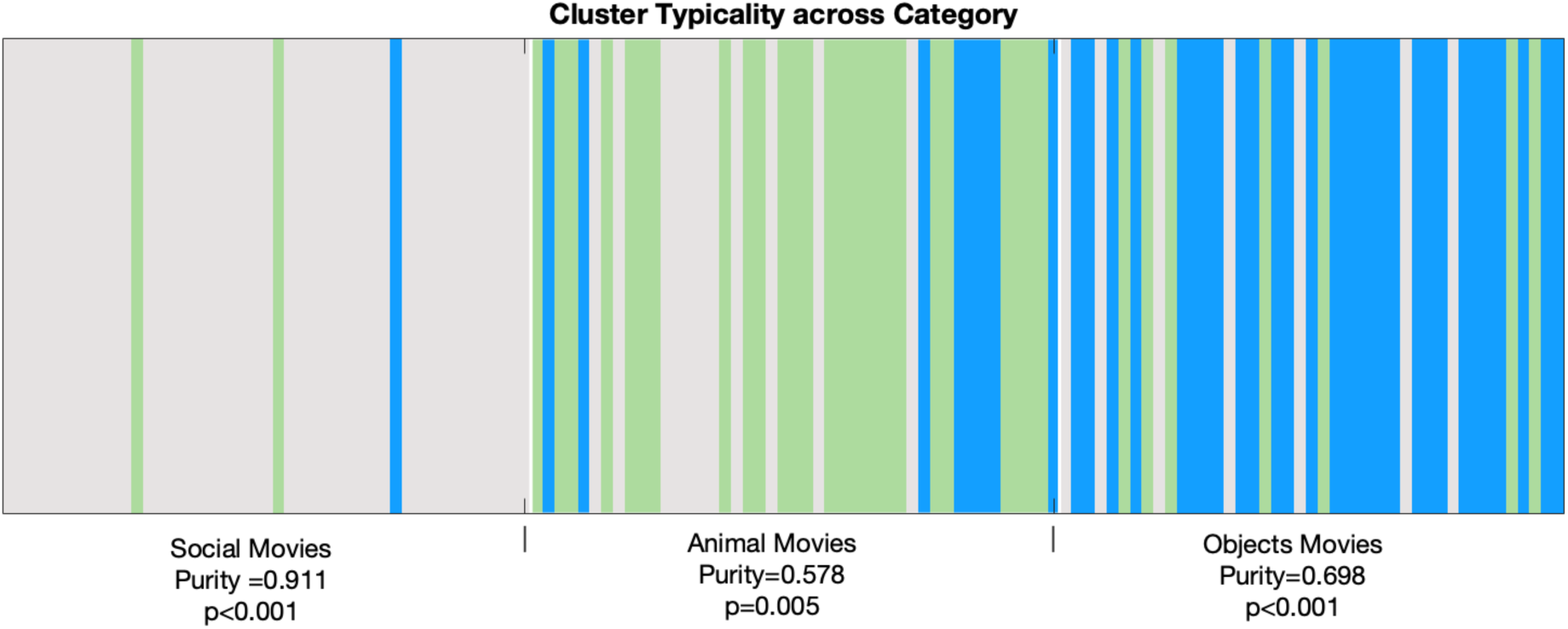
Clustering analysis of behavioral gaze typicality across movie categories. Hierarchical clustering of gaze typicality across all valid movies. Each vertical bar represents a movie, colored according to its assigned cluster. Movies are ordered by category, with animal, social, and object movies separated by the white lines. Cluster significance was assessed using permutation testing, and the observed purity exceeded the distribution obtained under random label assignments.

**Supplementary Figure S2.**
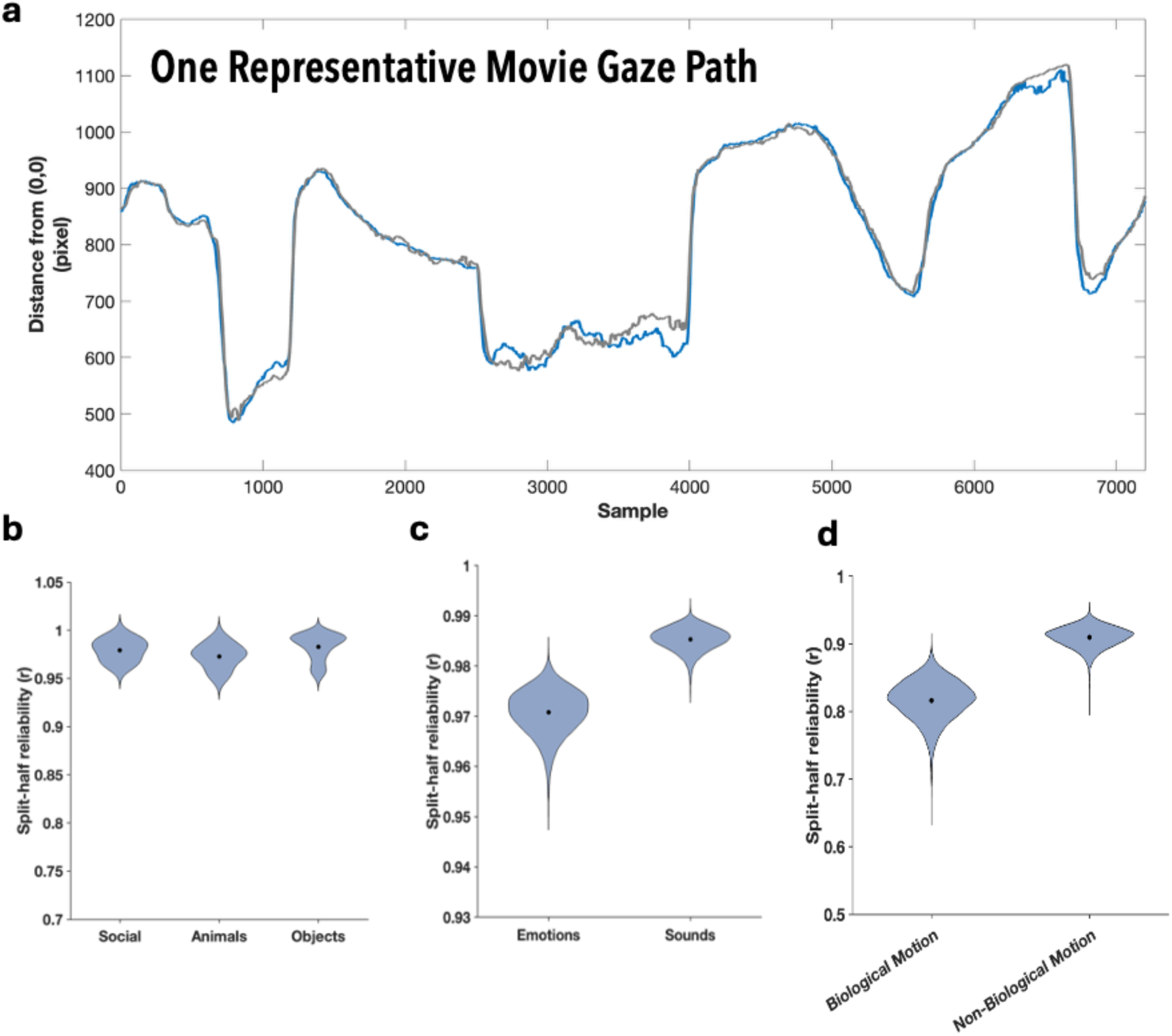
Split-half reliability of behavioral consensus across tasks. **(a**) Representative example of split-half reliability for gaze behavior during movie viewing. Participants were randomly divided into two independent groups, and the group-average gaze trajectory (distance from the upper-left corner of the screen) was computed separately for each group. The two independently derived consensus trajectories closely overlapped throughout the movie, illustrating the high reliability of the group-average gaze pattern**. (b)** Distribution of split-half reliability estimates across movies within each gaze category (Social, Animal, and Object movies), demonstrating consistently high reliability for the group consensus in all stimulus categories. **(c)** Split-half reliability of consensus ratings in the auditory task. Emotion-intensity and acoustic-intensity ratings both exhibited high reliability across repeated random split-half analyses, indicating robust agreement among participants. **(d)** Split-half reliability of prediction accuracy in the motion prediction task. Both biological and non-biological motion prediction showed high within-category reliability, demonstrating stable individual performance across independent subsets of stimuli. Black dots indicate the mean split-half correlation in each condition; violin plots represent the distribution of 10,000 random split-half iterations.

**Supplementary Figure S3.**
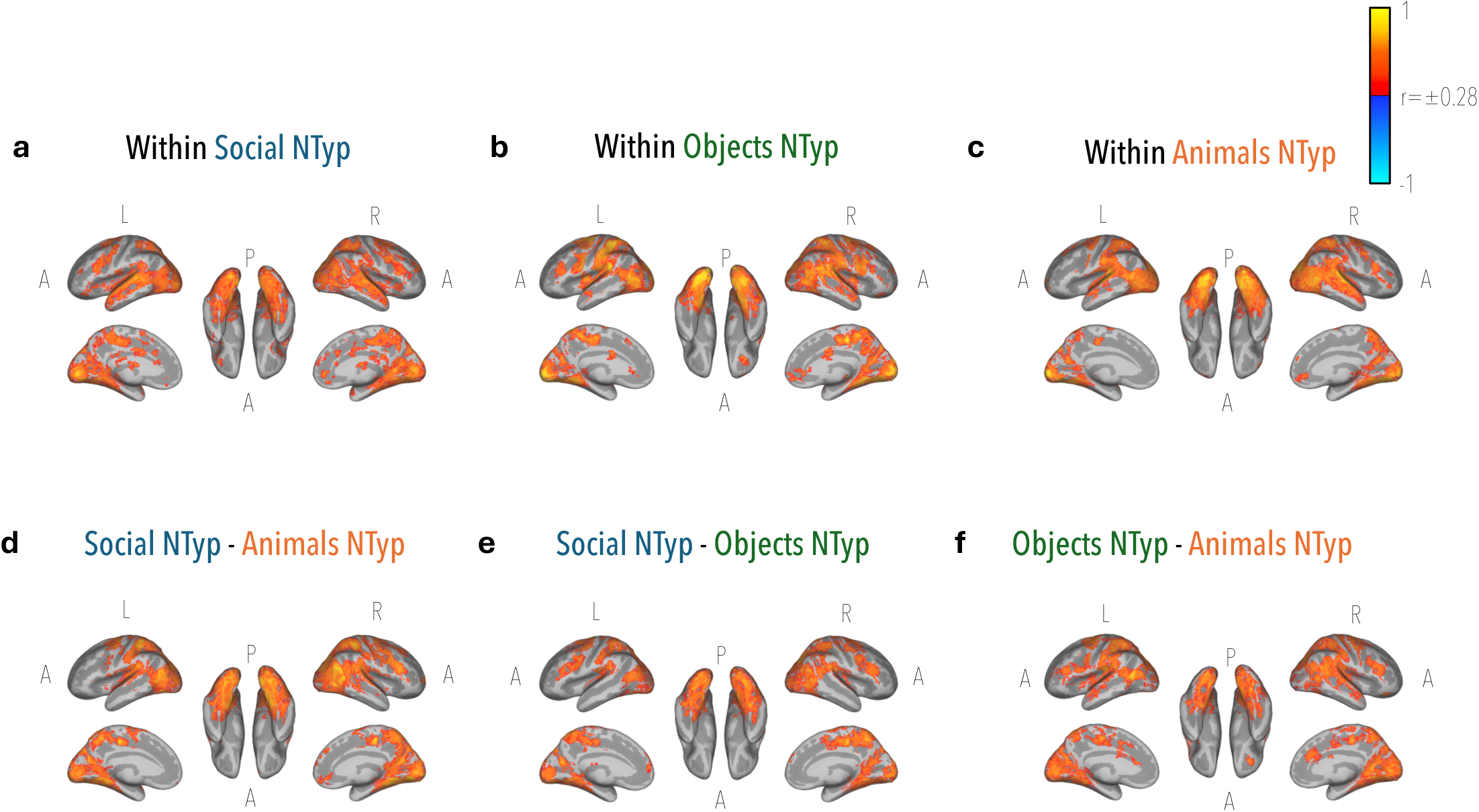
Stability of neural typicality within and across movie categories. Split-half reliability maps computed independently for social **(a)**, animal **(b)**, and object **(c)** movies demonstrate robust temporal stability of neural typicality within each movie category. Voxelwise comparisons of neural typicality across movie categories are shown for social versus animal **(d)**, social versus object **(e)**, and object versus animal **(f)**, demonstrating that stable neural typicality patterns are largely shared across naturalistic stimuli despite differences in movie content. Warm colors indicate positive correlations, and cool colors indicate negative correlations. Maps display only voxels surviving permutation-based cluster correction for multiple comparisons (cluster-corrected p < 0.05). Each analysis was thresholded using its own voxelwise and cluster-size thresholds, and only surviving clusters are shown.

**Supplementary Figure S4.**
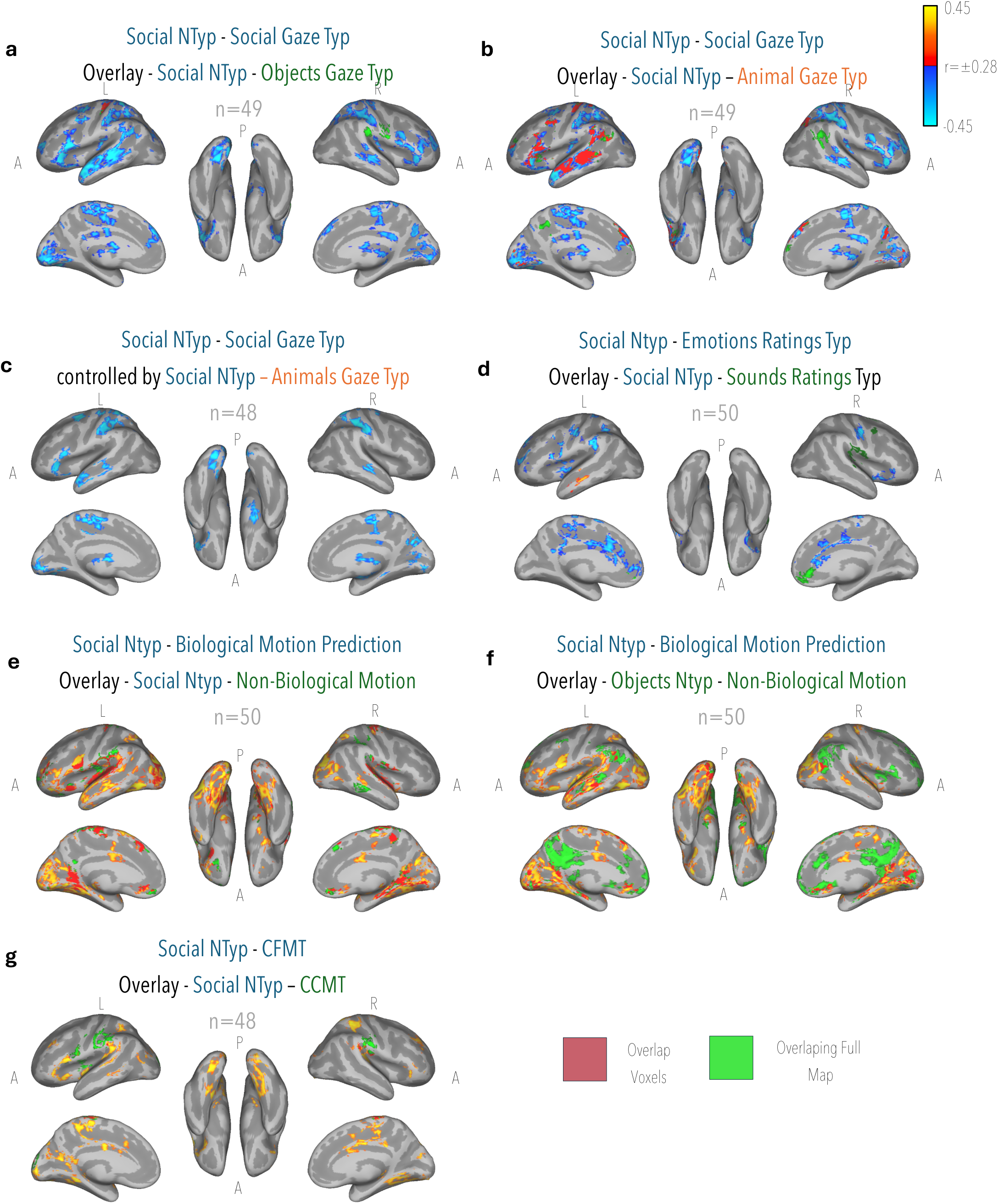
Correlations of neural typicality across movie categories. **(a)** Brain regions in which the association between social neural typicality and social movie viewing gaze typicality overlapped the association between social neural typicality and object movie viewing gaze typicality. **(b)** Brain regions in which the association between social neural typicality and social movie viewing gaze typicality overlapped the association between social neural typicality and animals movie viewing gaze typicality. **(c)** Brain regions in which the association between social neural typicality and social movie viewing gaze typicality remained significant after controlling for animals movie viewing gaze typicality. **(d)** Brain regions in which the association between social neural typicality and emotion rating typicality overlapped the association between social neural typicality and general sound ratings typicality. **(e)** Brain regions in which the association between social neural typicality and biological motion prediction overlapped the association between social neural typicality and non-biological motion prediction. **(f)** Brain regions in which the association between social neural typicality and biological motion prediction overlapped the association between objects neural typicality and non-biological motion prediction. **(g)** Brain regions in which the association between social neural typicality and CFMT overlapped the association between social neural typicality and CCMT.

**Supplementary Figure S5.**
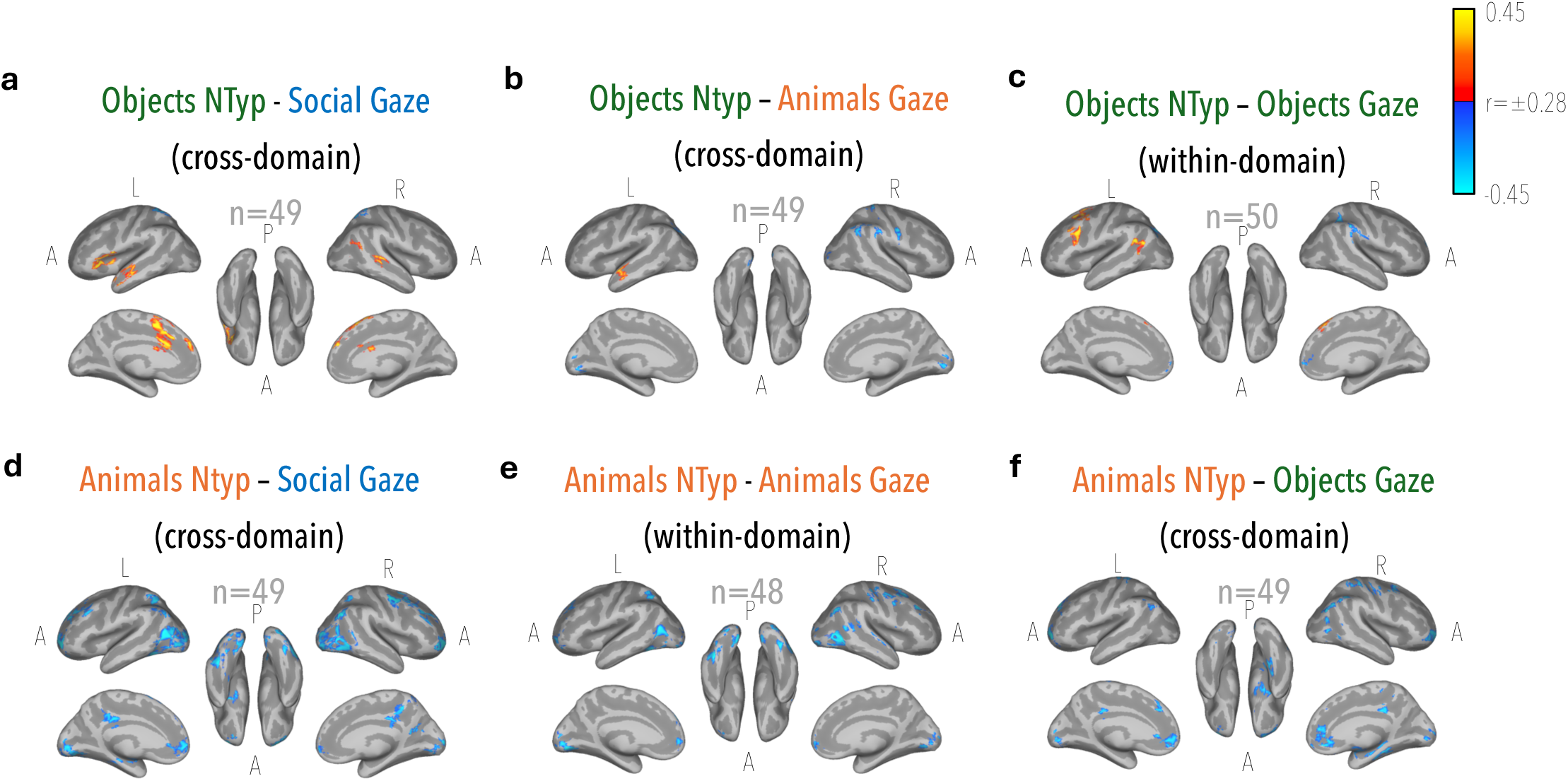
Voxelwise correlation maps showing the relationship between neural typicality (NTyp) during objects movie viewing and gaze typicality derived from social **(a)**, animal **(b)**, and object **(c)** movies, and voxelwise correlation maps showing the relationship between neural typicality (NTyp) during animals movie viewing and gaze typicality derived from social **(d)**, animal **(e)**, and object **(f)** movies.

**Supplementary Figure S6.**
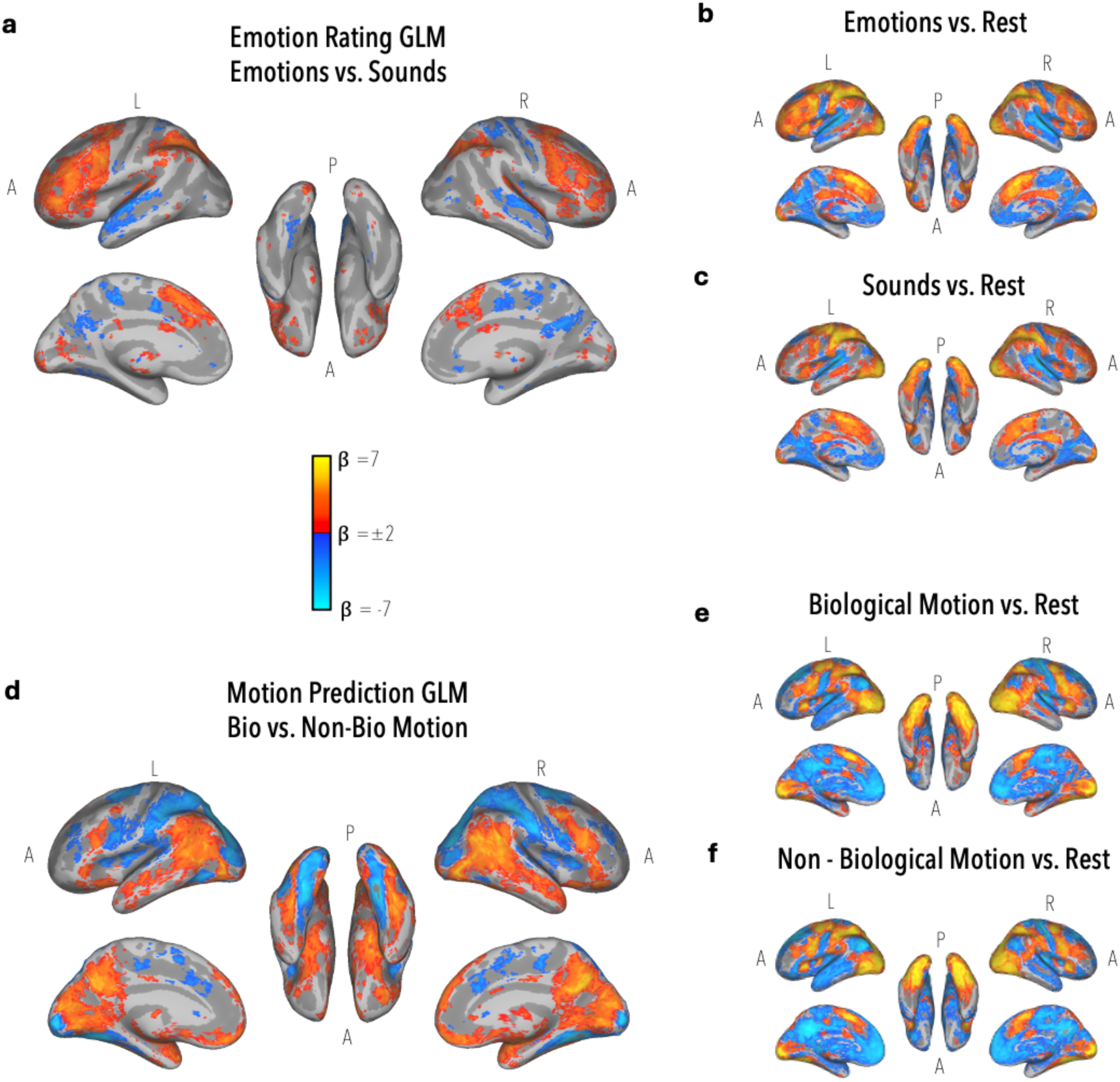
Whole-brain activation maps for the behavioral GLM tasks. Whole-brain GLM activation maps for the emotion-rating task **(a-c)** and the motion-prediction task (d-f). **(a)** Contrast between emotion ratings and sound ratings (Emotions vs. Sounds). **(b)** Brain regions associated with emotion ratings relative to the implicit baseline (Emotions vs. Rest). **(c)** Brain regions associated with sound ratings relative to the implicit baseline (Sounds vs. Rest). **(d)** Contrast between biological and non-biological motion prediction (Biological Motion vs. Non-Biological Motion). **(e)** Brain regions associated with biological motion prediction relative to the implicit baseline (Biological Motion vs. Rest). **(f)** Brain regions associated with non-biological motion prediction relative to the implicit baseline (Non-Biological Motion vs. Rest). Warm colors indicate positive β values, and cool colors indicate negative β values. Maps display only voxels surviving cluster-level correction for multiple comparisons (cluster-corrected *p* < 0.05). Each analysis was thresholded using its own voxelwise and cluster-size thresholds, and only surviving clusters are shown.

**Supplementary figure S7.**
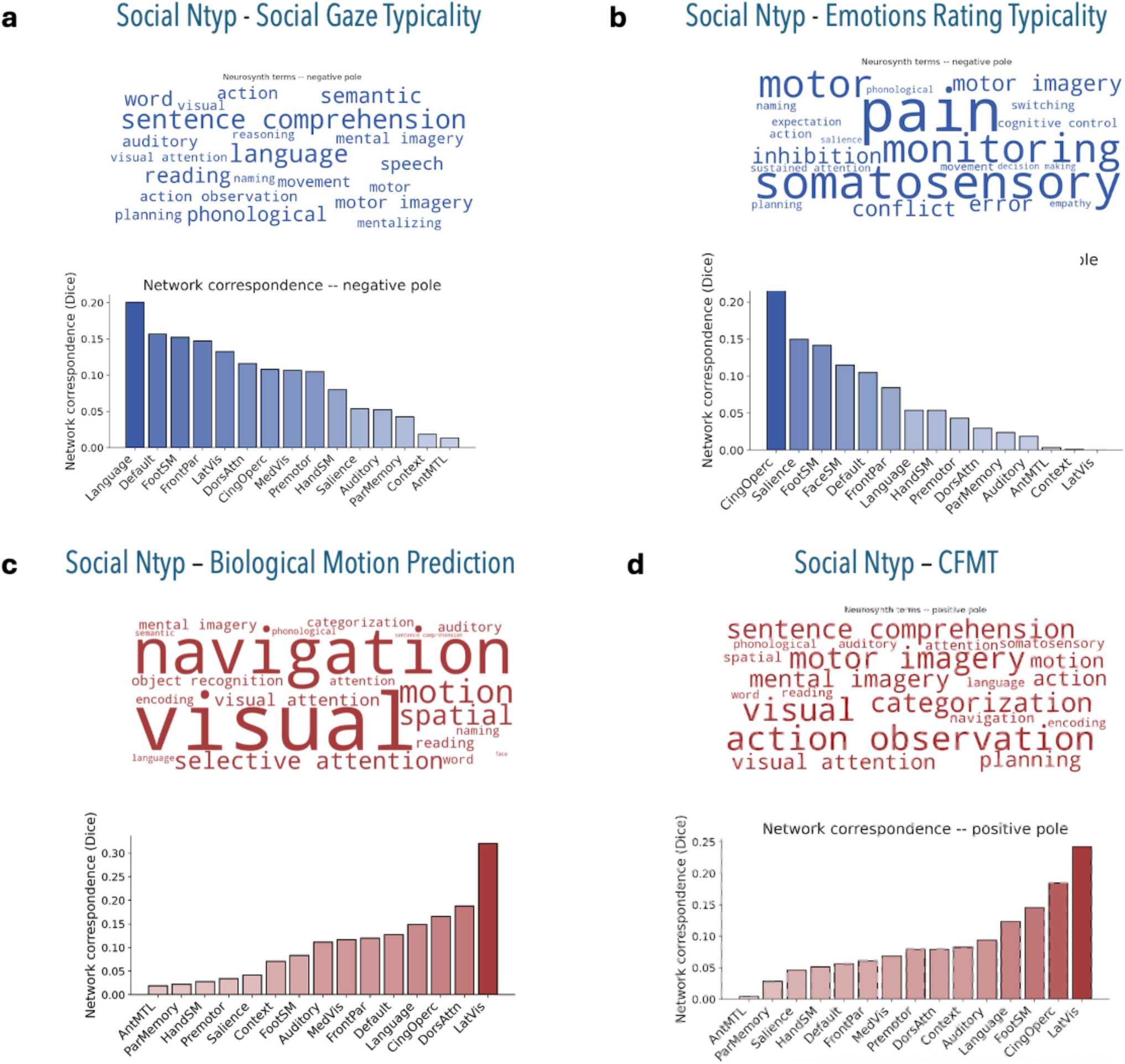
Semantic decoding and network correspondence index analysis for social neurotypicality and behavior maps. Each panel in the figure displays semantic terms associated with a given map (top), based on Neurosynth correlation Decoder as implemented in NiMare Python package (v0.20; Salo et al., 2023), and network correspondence index (Kong et al., 2025) based on Gordon 17 network atlas (Gordon et al., 2017). **(a)** social neural typicality correlation to social gaze typicality (Figure 4a); **(b)** social neural typicality correlation to emotions rating typicality (Figure 4d); **(c)** social neural typicality correlation to biological motion prediction (Figure 5a); **(d)** social neural typicality correlation to CFMT.

